# Generating and testing hypotheses about the fossil record of insect herbivory with a theoretical ecospace

**DOI:** 10.1101/2021.07.16.452692

**Authors:** Sandra R. Schachat, Jonathan L. Payne, C. Kevin Boyce, Conrad C. Labandeira

## Abstract

A typical fossil flora examined for insect herbivory contains a few hundred leaves and a dozen or two insect damage types. Paleontologists employ a wide variety of metrics to assess differences in herbivory among assemblages: damage type diversity, intensity (the proportion of leaves, or of leaf surface area, with insect damage), the evenness of diversity, and comparisons of the evenness and diversity of the flora to the evenness and diversity of damage types. Although the number of metrics calculated is quite large, given the amount of data that is usually available, the study of insect herbivory in the fossil record still lacks a quantitative framework that can be used to distinguish among different causes of increased insect herbivory and to generate null hypotheses of the magnitude of changes in insect herbivory over time. Moreover, estimates of damage type diversity, the most common metric, are generated with inconsistent sampling standardization routines. Here we demonstrate that coverage-based rarefaction yields valid, reliable estimates of damage type diversity that are robust to differences among floral assemblages in the number of leaves examined, average leaf surface area, and the inclusion of plant organs other than leaves such as seeds and axes. We outline the potential of a theoretical ecospace that combines various metrics to distinguish between potential causes of increased herbivory. We close with a discussion of the most appropriate uses of a theoretical ecospace for insect herbivory, with the overlapping damage type diversities of Paleozoic gymnosperms and Cenozoic angiosperms as a brief case study.

## 1 Introduction

In recent years, the number of fossil plant assemblages examined for insect herbivory has increased markedly. The wealth of available data has already been used to inform a variety of biotic and abiotic phenomena (Smith, 2008; Carvalho et al., 2014; Labandeira and Currano, 2013), but raises the question of how to compare the patterns of insect herbivory observed on different host plants or in different assemblages.

An increase in herbivory in deep time can occur in response to various environmental and evolutionary phenomena, demonstrating the need for analytical techniques that can be used to distinguish among them. Two explanations that are commonly invoked as causes of increased insect herbivory are insect and plant evolution, which lead to an expanded suite of feeding behaviors (Labandeira, 2006; Martinez et al., 2019; Wagner et al., 2015), and the nutrient dilution hypothesis, in which a sudden increase in atmospheric *p*CO_2_ increases the carbon-to-nitrogen ratio in many plant tissues, increasing the amount of leaf area that each insect must consume in order to ingest a given amount of protein (Bazzaz, 1990). The techniques currently used in paleontological studies do not distinguish among these disparate causes of increased herbivory.

A standardized method for comparing insect herbivory would allow the use of published data to generate null expectations for findings at new localities. In addition, a standardized method would facilitate the differentiation of these and other causes of increased insect herbivory in cases where the distinction is not so clear.

### 1.1 The end-triassic as a hypothetical case study

The end-Triassic extinction event exemplifies the potential utility of statistical methods with the capacity to generate null expectations and disentangle the various potential causes of fluctuations in herbivory. Few latest Triassic floras have been examined for insect herbivory (Ghosh et al., 2015) and, of the geologic periods that contain more than five described insect fossils, the Jurassic is the least studied in this context (McLoughlin et al., 2015; Ding et al., 2015; Pinheiro et al., 2016; Na et al., 2018; Santos et al., 2021).

One could generate any number of predictions about changes in insect herbivory across the end-Triassic event. Patterns of insect herbivory may have remained constant because it is widely agreed that insects did not suffer major losses at the Triassic/Jurassic boundary (Dmitriev and Zherikin, 1988; Labandeira and Sepkoski, 1993; Jarzembowski and Ross, 1996). Insect herbivory may have decreased because plant communities do appear to have endured noticeable turnover and losses across this extinction event (Belcher et al., 2010; Li et al., 2020; Lucas, 2021; McElwain and Punyasena, 2007). The *p*CO_2_ spike associated with the end-Triassic event (Knobbe and Schaller, 2017) complicates matters further. Whether or not an increase in *p*CO_2_ led to an increase in plant biomass and a corresponding dilution of nutrients such as nitrogen (Mattson, 1980) depends greatly on interacting environmental parameters (Shaw et al., 2002; McMurtrie et al., 2008; Reich et al., 2014). Nutrient dilution has very rarely been directly examined in the plant clades that were present in Triassic and Jurassic ecosystems; this phenomenon has been studied almost exclusively in angiosperms, which had not yet evolved at the time of the end-Triassic event (Bazzaz, 1990; Boyce and Zwieniecki, 2012; Ramírez-Barahona et al., 2020).

If data were available, a comparison of insect herbivory levels immediately before and after the end-Triassic event would be hampered by the lack of available statistical techniques. The first obstacle would be the lack of a null, or baseline, prediction of the magnitude of change in insect herbivory that would occur from the Late Triassic to Early Jurassic in the absence of a major environmental or evolutionary event. How much variation in insect herbivory is best attributed to statistical noise? How much is best attributed to the passage of time rather than an external trigger? After these sources of variation are taken into account, how much variation remains? The fern- and gymnosperm-dominated Permian, Triassic, and Cretaceous floras that have already been examined for insect herbivory provide an opportunity to generate a null prediction and quantify the uncertainty surrounding it. What is needed is a comparative framework to generate this null prediction.

The second obstacle would be the lack of a comparative framework for disentangling the biotic and abiotic causes of fluctuations in insect herbivory. The environmental perturbation most thoroughly examined in existing studies of insect herbivory, the Paleocene–Eocene Thermal Maximum, or PETM (Wilf and Labandeira, 1999; Currano et al., 2008, 2016), began and ended far too quickly for much plant or insect evolution to have occurred (Zeebe and Lourens, 2019). Many other environmental perturbations, such as the increase in *p*CO_2_ at the end-Triassic, which occurred in multiple pulses (Ruhl and Kürschner, 2011), span a long enough interval that abiotic and biotic factors can be confounded.

### 1.2 Theoretical ecospaces in paleontology

Morphospaces are a useful tool for quantifying change over time. The axes of empirical morphospaces, constructed with techniques such as principal component analysis, change with the addition of new data; in contrast, the axes of theoretical morphospaces remain unchanged as new data are added (McGhee, 2006). Morphospaces can be multidimensional (Raup, 1967; Lohman et al., 2017), can consist of various two-dimensional comparisons (Wilson and Knoll, 2010), or, with sufficiently clear and specific hypotheses, require only two dimensions (Raup, 1967; Gerber, 2017; Balisi and Van Valkenburgh, 2020).

Ecospaces extend the logic of empirical and theoretical morphospaces to ecological data. The canonical use of ecospace in paleontology applies to the marine realm (Valentine, 1969; Bambach, 1983), with an updated version now forming the foundation of many quantitative studies (Bush et al., 2007; Wiedl et al., 2013; Knope et al., 2015; Mondal and Harries, 2016; Frey et al., 2018; Laing et al., 2019). The three-dimensional ecospace as updated by Bush et al. (2007) has also been modified for the sedimentary ichnological study of terrestrialization (Minter et al., 2017) and for the study of terrestrial vertebrates (Chen et al., 2019). The primary difference between the two ecospace formulations currently used in studies of marine animals is the number of characters and character states (Novack-Gottshall, 2007; Bush and Novack-Gottshall, 2012). Of note, both theoretical ecospaces for marine animals use qualitative character states (Novack-Gottshall, 2007; Bush et al., 2007).

## 2 A theoretical ecospace for insect herbivory in the fossil record

Studies of insect herbivory typically use at least one of three common metrics. The diversity of insect damage types (Labandeira et al., 2007) measures the richness of herbivory. The percentage of leaf area removed by herbivores (known as the herbivory index) and the percentage of leaf specimens with feeding damage both measure the intensity of herbivory; however, the latter is highly sensitive to leaf size, behaves differently from the herbivory index (Smith, 2008), and thus is not recommended as an alternative to the former (Schachat et al., 2018). Although damage type diversity and the herbivory index are often discussed interchangeably, with an increase in either referred to as “more herbivory,” they measure fundamentally different aspects of herbivory (Figure 1).

**Figure 1:**
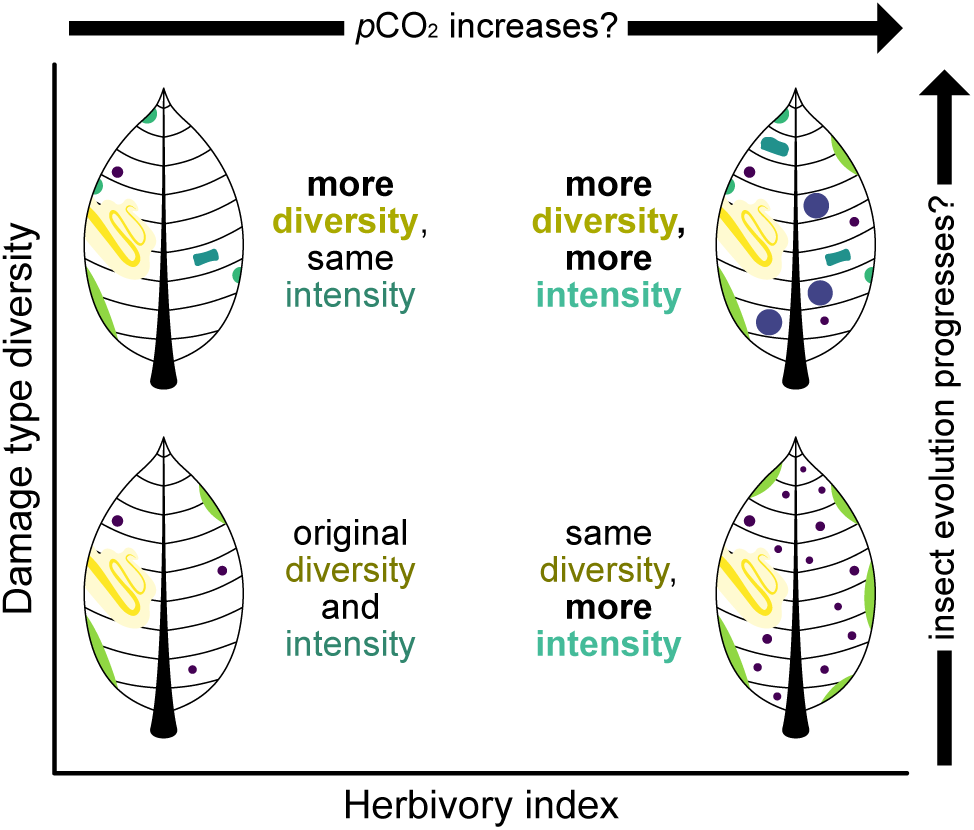
The difference between the diversity and intensity of insect herbivory.

The simplest theoretical ecospace for insect herbivory consists of damage type diversity plotted against the herbivory index. Although this ecospace has not been illustrated or analyzed in previous studies, it holds the potential to disentangle the various causes of increased herbivory.

### 2.1 The utility of an ecospace

The dilution of nitrogen in leaves, caused by increased plant biomass, in turn caused by increased *p*CO_2_, will increase the amount of leaf biomass that an herbivore will need to consume in order to ingest enough protein to satisfy its metabolic demands. In other words, an increase in *p*CO_2_ is predicted to cause an increase in the intensity of insect herbivory. The increased intensity of herbivory may well permit an increase in the observed diversity of herbivory because the more leaf mass is consumed, the more opportunities there are for the preservation of additional damage types that were present at low frequencies. However, this need not necessarily be the case. When the intensity of herbivory increases, the diversity of herbivory can remain constant or can even decrease.

In contrast, the expanding suite of insect feeding behaviors that developed from the Carboniferous through Permian increased the diversity of herbivory more measurably and consistently than it increased the intensity of herbivory. Whereas arthropod herbivory, probably caused by mites, has been found on Devonian liverworts (Labandeira et al., 2013), the oldest known insect herbivory in the fossil record consists only of external foliage feeding, with no accompanying piercing and sucking or specialized behaviors such as galling and mining (Iannuzzi and Labandeira, 2008). The size of the archetypal herbivorous arthropod and its feeding traces increased, but exhibited no more sophistication or diversity of feeding behavior than already seen with probable mite herbivory on Devonian liverworts. As another example, definitive evidence of leaf mining only appears in the earliest Mesozoic—tens of millions of years after insects began eating plants (Labandeira et al., 1994; Ding et al., 2014)—at a time when the diversity of gall-inducing arthropods also increased (Labandeira, 2021). An increase in the diversity of insect herbivory does not necessitate a corresponding increase in the intensity of herbivory: for example, some Permian gymnosperms have herbivory indices between 3.08% (Schachat et al., 2014) and 3.95% (Beck and Labandeira, 1998). These represent the twenty-sixth and thirty-second percentiles, respectively, of angiosperm leaf area removed per year by insect herbivores in extant ecosystems—slightly less than the mean value (Turcotte et al., 2014).

Thus, a bivariate theoretical ecospace can be used to differentiate among the expected impacts of nitrogen dilution versus insect and plant evolution as potential causes of increased insect herbivory. An increase in the diversity of herbivory that outpaces the intensity of herbivory is consistent with insect evolution but not with nitrogen dilution, and vice-versa.

The cyclical Holocene range expansions and contractions of eastern hemlock, *Tsuga canadensis* L., illustrate why the distinction between the diversity and intensity of herbivory matters (Labandeira, 2012). The decline of hemlock is believed to have been caused by either climate change or outbreaks of insect herbivores (Foster et al., 2006). Increased herbivory on subfossil hemlock leaves would support the hypothesis that insects drove eastern hemlock into decline, but only if an increase in the intensity of herbivory exceeds any increase in the diversity of herbivory. The intensity of herbivory would need to increase more strongly than the diversity of herbivory in order for the insect-outbreak hypothesis to be supported because, of the myriad insects known to feed on eastern hemlock (Buck et al., 2005; Dilling et al., 2007), only three—two moths and one conifer aphid—have been identified as possible culprits in the Holocene decline of this species (Filion et al., 2006; Simard et al., 2002; Oswald, 2016; Orwig and Foster, 1998; Labandeira, 2012). Moreover, due to indirect interspecific competition among insect herbivores (Janzen, 1973; Price et al., 1980; van Veen et al., 2006; Kaplan and Denno, 2007), an outbreak of a single herbivore species would quite possibly decrease the total diversity of insect herbivory through suppression of other herbivores. Therefore, range contractions of eastern hemlock caused by insect herbivores would require an increase in the intensity, but not the diversity, of herbivory.

### 2.2 Estimating damage type diversity for an ecospace

Because differences in sampling completeness threaten to bias estimates of damage type diversity (Schachat et al., 2018), the act of comparing damage type diversity to the herbivory index is no simple task. Estimates of the herbivory index are largely robust to differences in sample size: as sampling intensity increases, confidence intervals become narrower but point estimates of the herbivory index do not change perceptibly because a mean is an unbiased estimator (Schachat et al., 2018). For damage type diversity, on the other hand, point estimates and the ranges of confidence intervals vary substantially with sampling intensity because a tally—in this case, a tally of the damage types observed—is a biased estimator that continues to increase as sampling progresses. This is illustrated by the Colwell Creek Pond and Mitchell Creek Flats assemblages from the Early Permian of Texas (Schachat et al., 2014, 2015) which overlap considerably in composition of their floral communities but vary in sampling completeness; the former contains over fifteen times as much broadleaf surface area and over five times as many broadleaf specimens as the latter (Figure S1). As detailed in the supplemental material, we found that the Chao1 estimator is unable to overcome the lack of sampling completeness in insect herbivory datasets. This estimator returns inaccurate and imprecise estimates of damage type diversity whether used alone (Chao, 1987) or in conjunction with rarefaction (Chao et al., 2014).

Coverage-based rarefaction provides an alternative to traditional rarefaction (Chao and Jost, 2012). In traditional, or size-based, rarefaction, curves are scaled by the number of leaves or the amount of leaf area examined. In coverage-based rarefaction, samples are standardized by sampling completeness as indicated by the slope of the rarefaction curve (Good, 1953; Jost, 2010). At the base of a rarefaction curve, sampling is incomplete; the slope of the rarefaction curve equals 1 and coverage equals 0. As sampling approaches completeness, the slope of the rarefaction curve reaches an asymptote—*i.e.*, a slope of 0—and coverage equals 1 (Figure 2). Thus, with coverage-based rarefaction, damage type datasets are rarefied not to a particular number of leaves but to a particular slope of the rarefaction curve. If a high proportion of damage types are observed on only one specimen, this indicates that coverage is relatively low. In contrast, if a low proportion of damage types are observed on only one specimen, coverage is relatively high. Coverage-based rarefaction is performed by removing the rarest damage types from the dataset until coverage is reduced to a predetermined threshold such as 0.8 (Figure 3).

**Figure 2:**
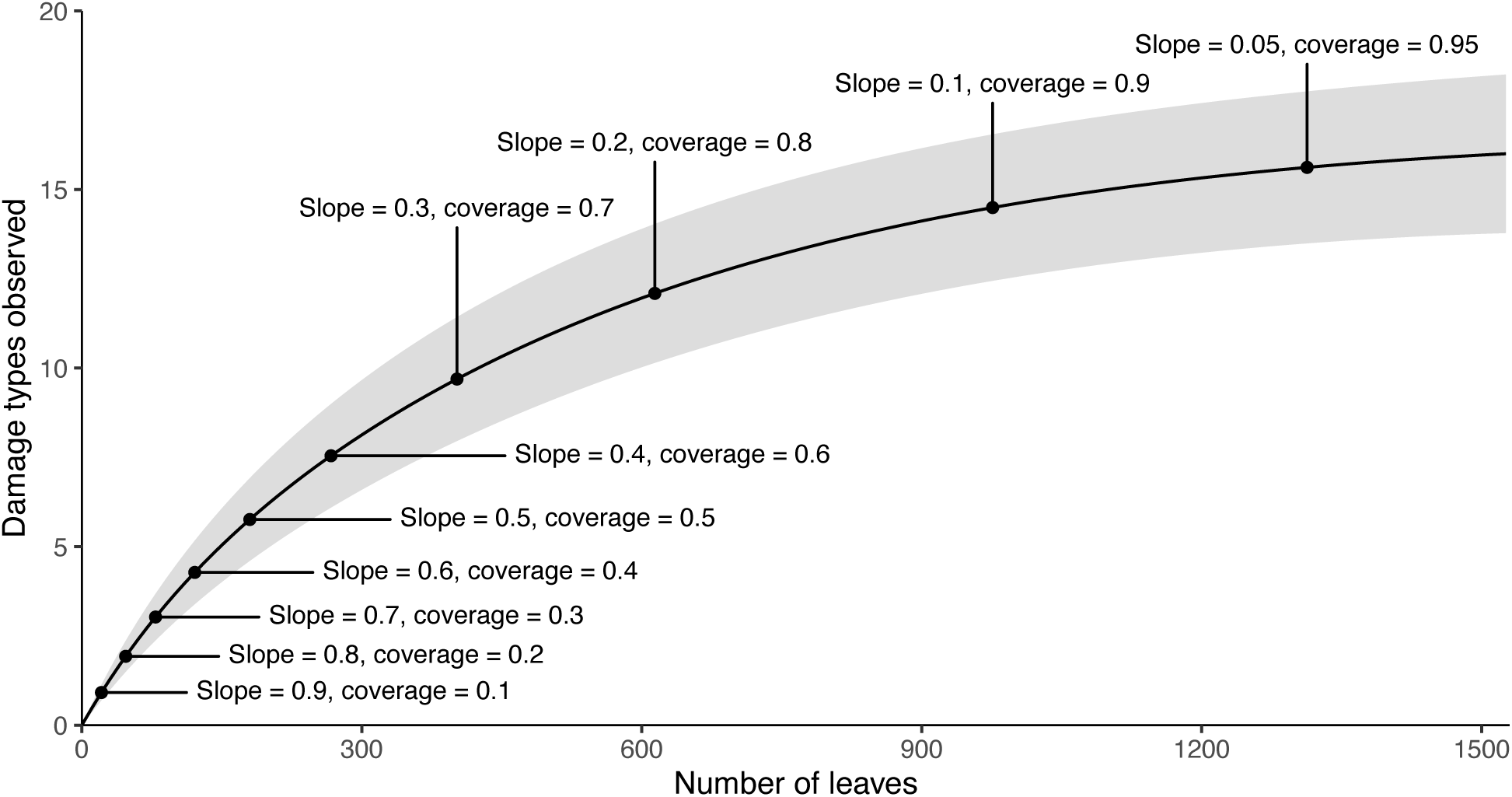
An illustration of the relationship between size-based and coverage-based rarefaction, showing that sample coverage is determined by subtracting the slope of the rarefaction curve from 1. These data are from the Hlatimbe Valley 213 assemblage (Labandeira et al., 2018), which was selected for inclusion here because it is also featured in Figure 3.

**Figure 3:**
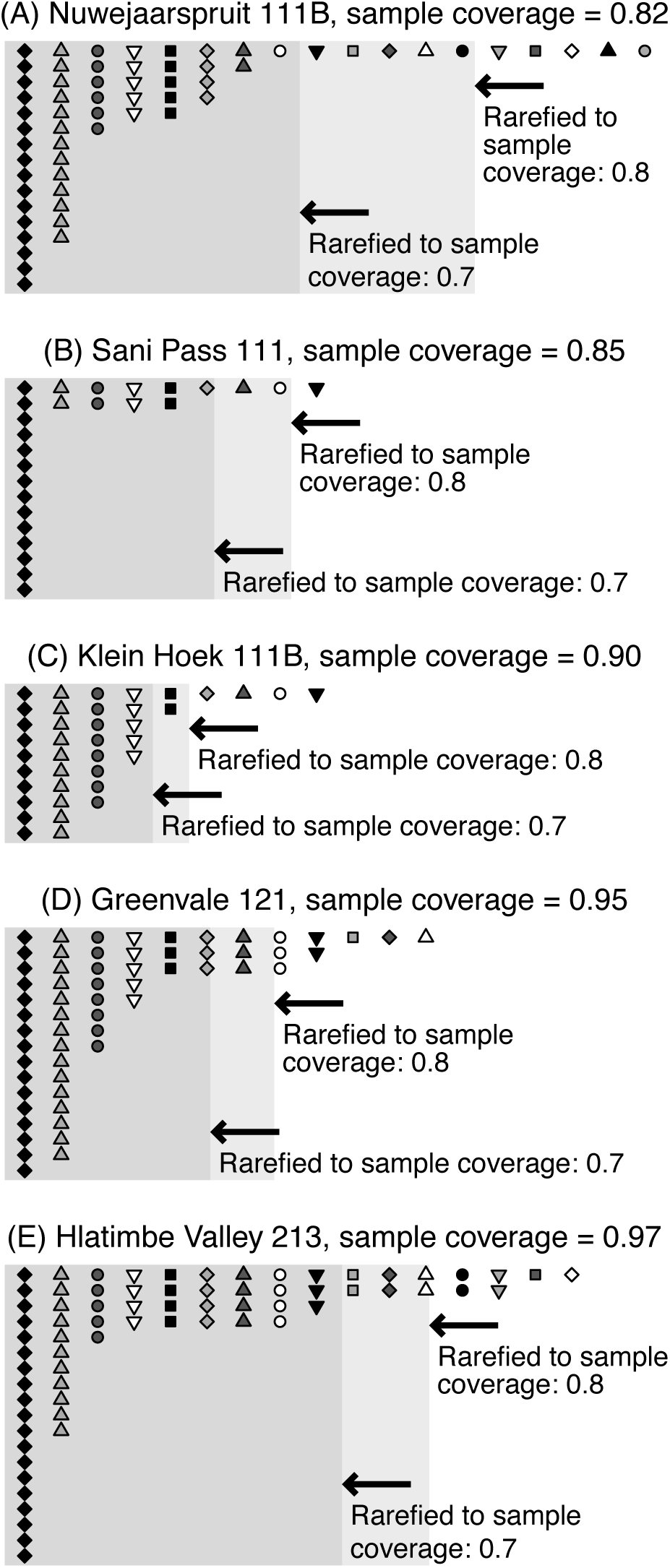
An illustration of the concepts of sample coverage and of rarefaction by sample coverage. The five assemblages illustrated here were examined by Labandeira et al. (2018). For each assemblage, the sample coverage for the complete dataset is listed in the title. The lightest gray boxes denote the damage type diversity at each assemblage when rarefied to a sample coverage of 0.8, and the darker gray boxes denote damage type diversity rarefied to a sample coverage of 0.7. The assemblages are organized from the lowest to highest levels of sample coverage. Each column of symbols represents a damage type, and the number of times each symbol is illustrated represents the number of specimens on which the damage type was observed. (A) has relatively low sample coverage because the majority of damage types are observed on only one specimen. In contrast, (E) has relatively high sample coverage because so few damage types are observed on only one specimen.

Because coverage-based rarefaction follows the replication principle (Chao and Jost, 2012) it provides an unbiased and consistent estimator of damage type diversity that is robust to sample size, leaf size, and fragmentation. In statistical parlance, an “unbiased estimator” is an estimator whose average expected value for a sample is equal to the true value in the population from which the sample was drawn. In other words, whether the dataset for a given fossil assemblage reaches a sample coverage of 0.85 or 0.99, the estimated damage type diversity will, on average, be the same when the dataset is rarefied down to a sample coverage of 0.8. A “consistent estimator” converges on the true population value as sample size becomes large. In other words, if one fossil assemblage reaches a sample coverage of 0.85 and another reaches a sample coverage of 0.99, the uncertainty surrounding damage type diversity rarefied to a sample coverage of 0.8 will be smaller for the latter, more completely-sampled assemblage. The assemblages listed in Tables 1 and 2 with over 7,000 leaves examined were randomly subsampled down to 1,000 and 2,000 leaves in a procedure that was iterated 1,000 times (Figure 4). Whereas accuracy and precision typically suffer when these assemblages are subsampled down to 1,000 or even 2,000 leaves before performing traditional, size-based rarefaction (Figure S3), the loss of precision and accuracy are minimal with coverage-based rarefaction (Figure 4). The loss of precision with coverage-based rarefaction is far less severe than with size-based rarefaction (Figure S3) or with the Chao1 estimator (Figure S2) and still allows the differentiation of damage type diversity among many assemblages. With coverage-based rarefaction, the most notable lack of overlap among the confidence intervals for the different sample sizes occurs for Clouston Farm (Figure 4). The confidence intervals overlap to a reassuring extent for half of the pre-angiosperm and all of the angiosperm assemblages (Figure 4).

**Figure 4:**
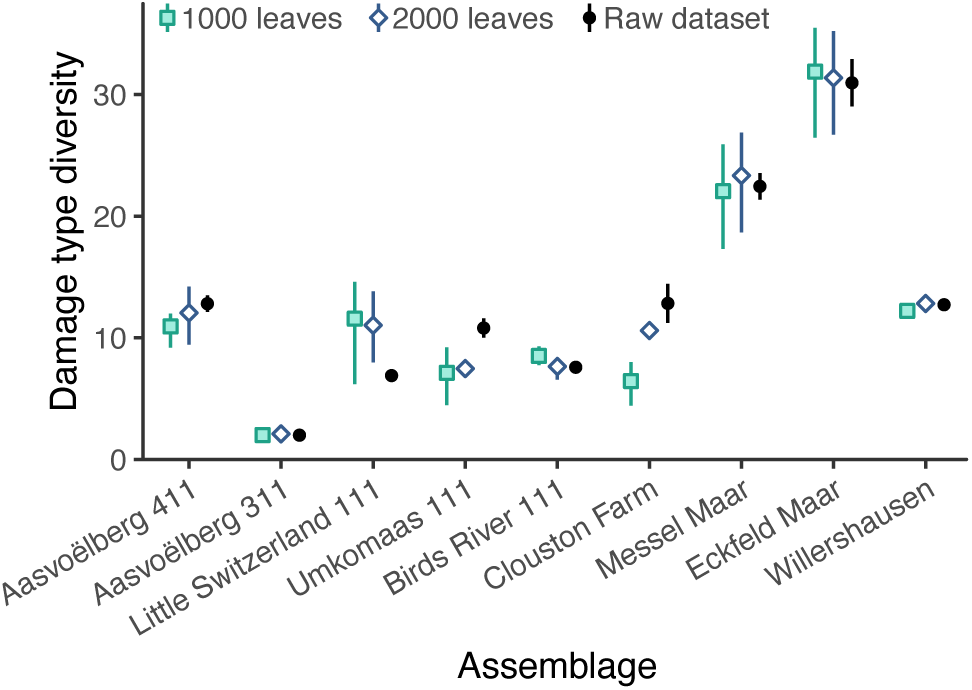
Precision and accuracy of estimates of damage type diversity generated with coverage-based rarefaction.

**Table 1:** The assemblages with over 1,000 broadleaf specimens from Labandeira et al. (2018), collected from the Molteno formation by Anderson and Anderson (1983, 1985, 1989, 2003, 2008, 2017), that were analyzed in this study. ^a^These assemblages do not have sufficient sample coverage to be rarefied to a coverage of 0.8.

| Assemblage | Leaves | DTs | Period | Age | Citation |
| --- | --- | --- | --- | --- | --- |
| Birds River 111 | 7,106 | 35 | Triassic | Carnian | Labandeira et al. (2018) |
| Greenvale 121 | 2,749 | 12 | Triassic | Carnian | Labandeira et al. (2018) |
| Boesmanshkoek 112 | 1,058 | 2 | Triassic | Carnian | Labandeira et al. (2018) |
| Cyphergat 111A | 5,499 | 26 | Triassic | Carnian | Labandeira et al. (2018) |
| Kannaskop 111 | 2,114 | 11 | Triassic | Carnian | Labandeira et al. (2018) |
| Telemachus Spruit 111 | 5,860 | 14 | Triassic | Carnian | Labandeira et al. (2018) |
| Kommandantskop 111 | 1,096 | 8 | Triassic | Carnian | Labandeira et al. (2018) |
| Vineyard 111 | 2,146 | 10 | Triassic | Carnian | Labandeira et al. (2018) |
| Elandspruit 111 | 1,054 | 10 | Triassic | Carnian | Labandeira et al. (2018) |
| Kraai River 311 | 1,213 | 6 | Triassic | Carnian | Labandeira et al. (2018) |
| Kraai River 111<sup>a</sup> | 1,780 | 8 | Triassic | Carnian | Labandeira et al. (2018) |
| Lutherskop 311 | 5,180 | 22 | Triassic | Carnian | Labandeira et al. (2018) |
| Waldeck 111 | 1,322 | 11 | Triassic | Carnian | Labandeira et al. (2018) |
| Konings Kroon 222<sup>a</sup> | 1,679 | 13 | Triassic | Carnian | Labandeira et al. (2018) |
| Konings Kroon 111A | 1,073 | 5 | Triassic | Carnian | Labandeira et al. (2018) |
| Peninsula 321 | 1,311 | 4 | Triassic | Carnian | Labandeira et al. (2018) |
| Peninsula 311 | 1,665 | 12 | Triassic | Carnian | Labandeira et al. (2018) |
| Peninsula 411 | 6,254 | 17 | Triassic | Carnian | Labandeira et al. (2018) |
| Klein Hoek 111B | 1,004 | 9 | Triassic | Carnian | Labandeira et al. (2018) |
| Klein Hoek 111C | 2,644 | 17 | Triassic | Carnian | Labandeira et al. (2018) |
| Kappokraal 111 | 1,453 | 27 | Triassic | Carnian | Labandeira et al. (2018) |
| Elandspruit 112A | 1,131 | 5 | Triassic | Carnian | Labandeira et al. (2018) |
| Nuwejaarspruit 111B | 1,582 | 18 | Triassic | Carnian | Labandeira et al. (2018) |
| Winnaarspruit 111 | 1,396 | 16 | Triassic | Carnian | Labandeira et al. (2018) |
| Morija 111B | 1,235 | 7 | Triassic | Carnian | Labandeira et al. (2018) |
| Makoaneng 111 | 1,206 | 6 | Triassic | Carnian | Labandeira et al. (2018) |
| Hlatimbe Valley 213 | 1,526 | 16 | Triassic | Carnian | Labandeira et al. (2018) |
| Umkomaas 111 | 9,828 | 36 | Triassic | Carnian | Labandeira et al. (2018) |
| Sani Pass 111 | 1,236 | 9 | Triassic | Carnian | Labandeira et al. (2018) |
| Qachasnek 111<sup>a</sup> | 2,093 | 7 | Triassic | Carnian | Labandeira et al. (2018) |
| Matatiele 111 | 3,667 | 23 | Triassic | Carnian | Labandeira et al. (2018) |
| Golden Gate 111 | 1,208 | 16 | Triassic | Carnian | Labandeira et al. (2018) |
| Little Switzerland 111 | 8,218 | 28 | Triassic | Carnian | Labandeira et al. (2018) |
| Aasvoëlberg 111 | 2,941 | 11 | Triassic | Carnian | Labandeira et al. (2018) |
| Aasvoëlberg 211 | 1,942 | 12 | Triassic | Carnian | Labandeira et al. (2018) |
| Aasvoëlberg 311 | 10,809 | 18 | Triassic | Carnian | Labandeira et al. (2018) |
| Aasvoëlberg 411 | 12,398 | 39 | Triassic | Carnian | Labandeira et al. (2018) |

**Table 2:**
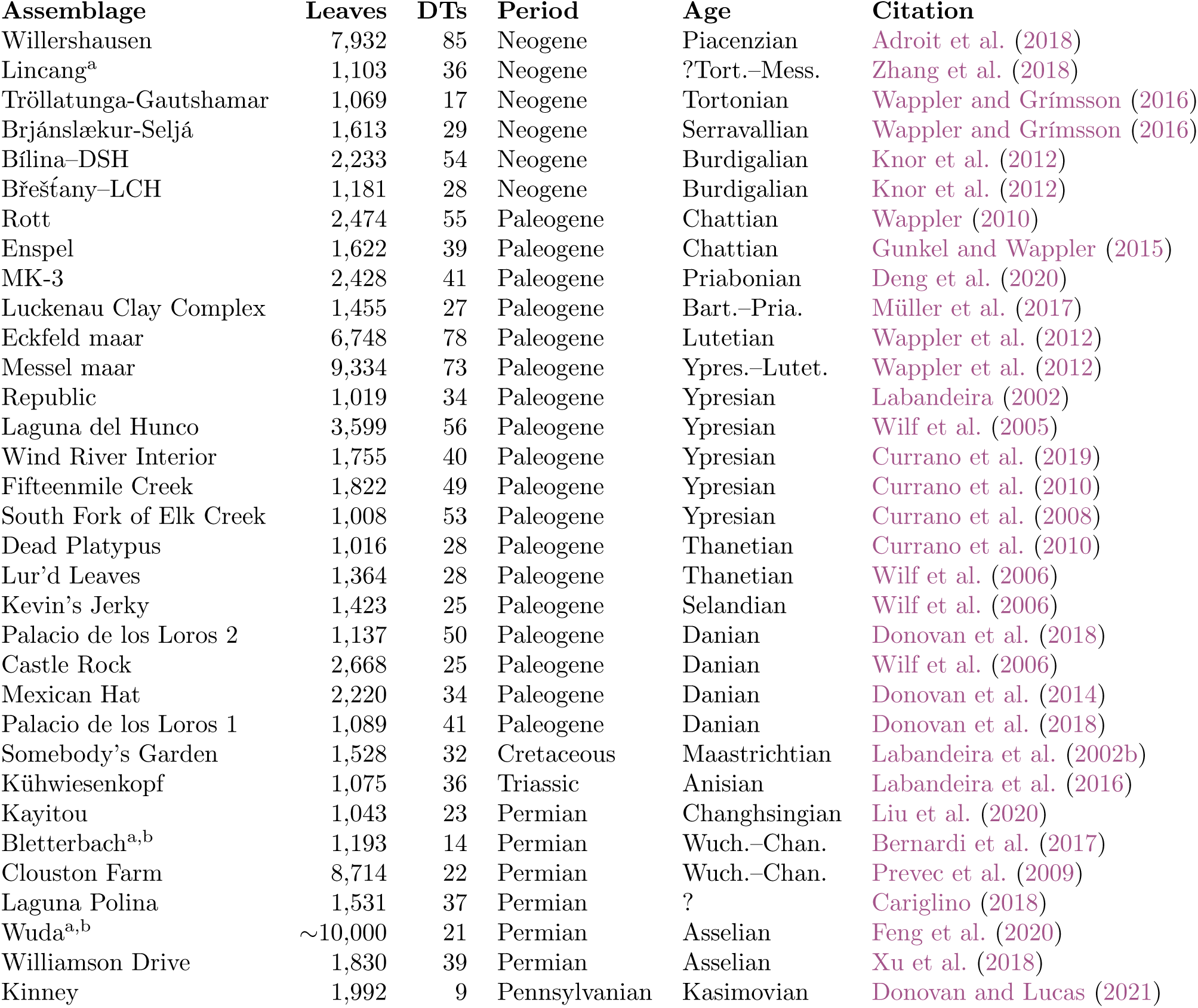
The published assemblages with over 1,000 broadleaf specimens from sources other than Labandeira et al. (2018) that were analyzed in this study. ^a^These assemblages are not included in the sensitivity analysis presented in Figure S2 because damage type data are not available for each individual specimen. ^b^These assemblages do not have sufficient sample coverage to be rarefied to a coverage of 0.8.

An additional benefit of coverage-based rarefaction is that, because samples are standardized by damage type completeness rather than number of leaves examined, this method is relatively robust to the variable inclusion of non-broadleaf specimens, from pine needles to axes to seeds, in the datasets for different assemblages. Some published comparisons of insect herbivory among assemblages include broadleaf specimens only, whereas others include all plant specimens. Some floral assemblages do not contain any seeds and other non-foliar specimens, and of those that do, the non-foliar specimens may or may not be included in published datasets. Within angiosperm-dominated floras, ferns, gymnosperms, and monocots may (Robledo et al., 2018; Giraldo et al., 2021) or may not (Azevedo Schmidt et al., 2019; Currano et al., 2019) be examined. Criteria for inclusion in comparative analyses can even vary within the same research group, with some workers using a wider definition of foliage that includes needles, liverworts, phyllids, photosynthetic wings of seeds, and even flattened horsetail axes (Prevec et al., 2009; Labandeira et al., 2018), and others using a narrower definition restricted to multi-veined broad leaves or leaves with a defined midvein (Schachat et al., 2014, 2015, 2018, 2020). (For additional information, see “criteria for inclusion of leaves and damage types” in the supplemental information.)

Because non-broadleaf specimens typically contain little or no evidence of insect herbivory, they contribute little or nothing to metrics of damage type sampling completeness and will therefore have minimal or no impact on estimates of damage type diversity at an assemblage. And, because some herbivory datasets code plants as morphotypes only, with no information on whether the morphotype represents a dicot or a conifer (Maccracken, 2020), these datasets cannot be analyzed with traditional size-based rarefaction. However, these datasets can indeed be analyzed with coverage-based rarefaction. Plant organs that contain little or no insect damage, such as axes and pine needles, contribute little or nothing to estimates of sample coverage, and thus their inclusion in coverage-based damage diversity estimates does not bias the estimates downward.

A remaining question is the confidence level that should be used in confidence intervals for rarefied damage type diversity. Schachat et al. (2018) used 84% confidence intervals, rather than 95%, because the comparison of two curves with 84% confidence intervals yields a Type I error rate below 5%. However, the confidence intervals generated with the iNEXT package rely in part on extrapolated diversity, which is shown above to be biased by sample size in the case of damage type diversity datasets. As a result, the confidence intervals generated by iNEXT are narrower than those generated with the iterative procedure of Schachat et al. (2018). Therefore, we recommend generating 95% confidence intervals with the iNEXT package for use in the ecospace.

#### 2.2.1 Considerations for assemblages with insufficient coverage

Just as size-based rarefied estimates of damage type diversity are conducted at a predetermined level of sampling, such as 400 leaves, coverage-based rarefied estimates of damage type diversity require a predetermined level of coverage, or sampling completeness. Here we rarefy to sample coverage 0.8. Coverage increases throughout the sampling process as a function of sample size, the evenness of damage types, the richness of damage types, and the frequency with which damage types occur. Accordingly, there is no predetermined number of leaves or amount of surface area at which an assemblage is assured to have sufficient coverage.

The impossibility of assessing all assemblages that have been sampled to a certain size, such as 400 leaves or 1000 cm^2^ of surface area, is perhaps the only disadvantage of using coverage-based rarefaction instead of size-based rarefaction. Five of the assemblages in Tables 1 and 2 have insufficient coverage for rarefaction to sampling completeness of 0.8: the Permian Wuda (Feng et al., 2020) and Bletterbach (Bernardi et al., 2017) assemblages, and the Triassic Qachasnek 111, Kraai River 111, and Konings Kroon 222 assemblages (Labandeira et al., 2018). (The Triassic Makoaneng 111 and Konings Kroon 111A assemblages both have a total sample coverage of 0.79. We extrapolated the damage type diversity at a sample coverage of 0.8 for these two asseblages, but do not recommend extrapolating diversity estimates for assemblages with coverage below 0.79, as discussed in the supplemental material.) All of these contain little damage. Of note, the Boesmanshkoek 112 assemblage from the same Triassic study contains only two damage types, but these occur frequently enough that the assemblage has sufficient coverage. Thirty-nine of the 44 assemblages with fewer than 1,000 leaves listed in Table 3 have sufficient coverage as well. Of the ten assemblages with coverage below 0.79, eight have values between 0.45 and 0.69, and the remaining two assemblages have values of 0.70 and 0.72. Rarefying to a sample coverage of 0.7 would decrease the power to detect differences among assemblages while providing a negligible increase in the number of assemblages included.

**Table 3:**
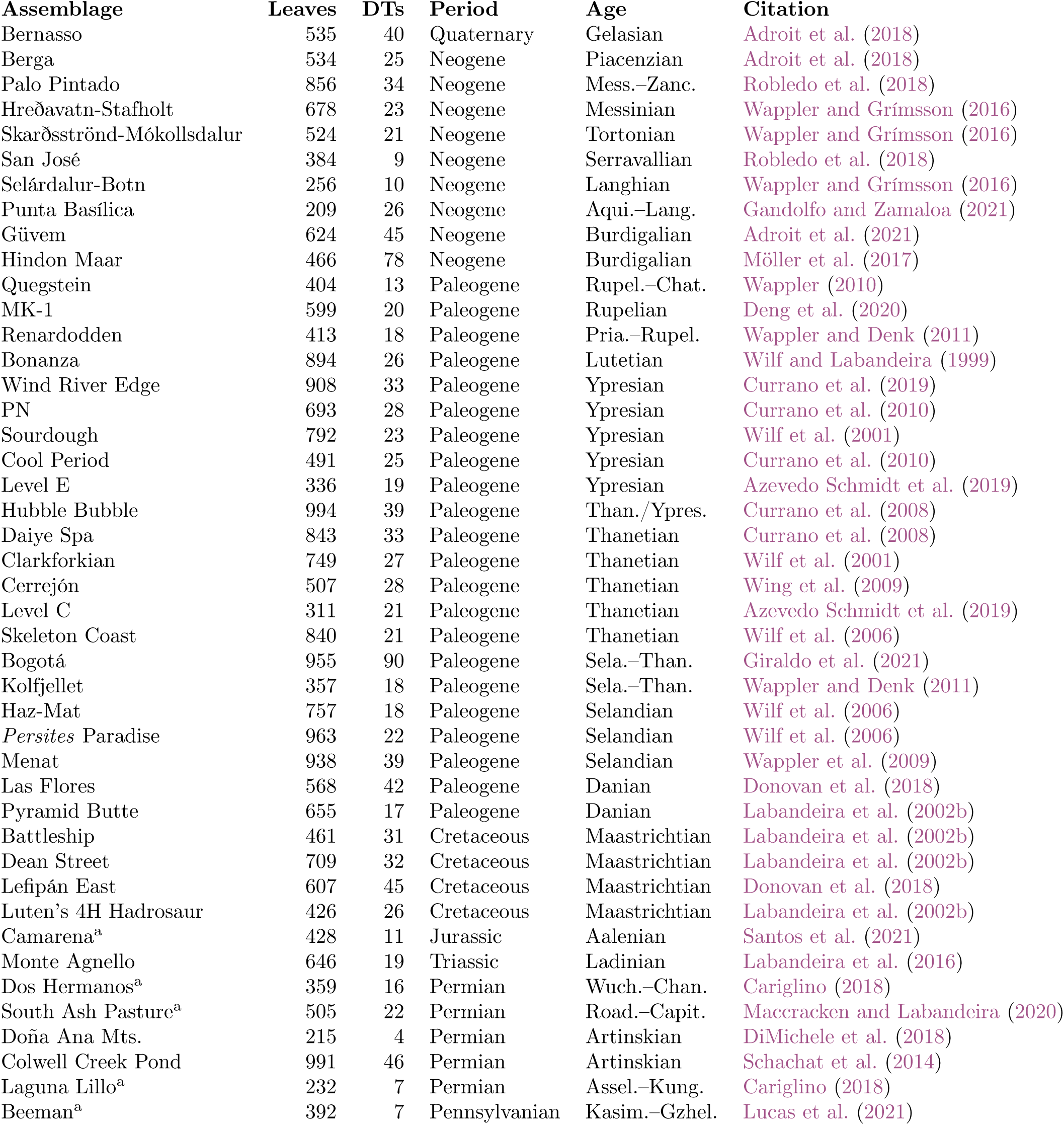
The assemblages analyzed in this study with 200–1,000 broadleaf specimens. ^a^These assemblages do not have sufficient sample coverage to be rarefied to a coverage of 0.8.

The best way to handle assemblages with insufficient coverage depends on the number of leaves examined and the temporal scope of the study. In the case of assemblages with only a few hundred leaves, estimates of damage type diversity comparable to those generated with coverage-based simply cannot be calculated. For studies such as that of Labandeira et al. (2002b) with a limited temporal scope and a large quantity of assemblages, many of which contain small numbers of leaves, the exclusion of assemblages that do not reach sufficient sample coverage can decrease the power of the study to detect changes in herbivory through time by excluding assemblages whose low sample coverage may be due to insufficient sampling or a true scarcity of damage types. In cases such as this, size-based rarefaction curves scaled by the amount of surface area sampled rather than the number of leaves sampled (Schachat et al., 2018) are the most appropriate choice. These size-based estimates of damage type diversity cannot be incorporated into meta-analyses that use coverage-based rarefaction. We conducted simulations to evaluate extrapolated estimates of damage type diversity in cases when the raw dataset does not reach a sample coverage of 0.8, and found that extrapolated estimates are not reliable and are often invalid (Figure S4).

### 2.3 Evaluating other potential dimensions of an ecospace for herbivory

Whereas the intensity and diversity of insect damage are the two most commonly studied aspects of insect herbivory and are the two most obvious choices for dimensions of an ecospace, additional dimensions bear consideration. Examples discussed here include the proportion of feeding occurrences belonging to external foliage feeding and to piercing and sucking; damage type evenness; damage type evenness compared to floral evenness; floral diversity; damage type diversity compared to floral diversity; and the prevalence of each functional feeding group, best quantified through the amount of herbivorized surface area. Feeding occurrences are defined here as the number of times that a damage type occurs on an individual plant specimen; these data are rarely collected in studies of fossil herbivory (Robledo et al., 2018; Ma et al., 2020). The first additional dimension is the prevalence of external foliage feeding, a functional feeding group containing generalist modes of herbivory in which an insect typically chews on a leaf. The proportion of feeding occurrences belonging to external foliage feeding—as opposed to piercing and sucking or specialized herbivory such as mining and galling—is perhaps the metric with the greatest potential to distinguish between insect evolution and nutrient dilution, the two most common explanations of increased herbivory invoked by paleontologists.

When nutrient dilution occurs, the prevalence of piercing-and-sucking feeding damage—whether measured as number of damage types, percentage of leaf area damaged, or number of feeding occurrences—will likely remain constant. This is because piercing-and-sucking insects often feed by puncturing individual phloem cells (Will et al., 2013). Whereas the number of phloem cells may increase in response to nutrient dilution, the content of each cell and the pressure within each cell will most likely remain the same. Therefore, piercing-and-sucking insects that feed on phloem will not need to feed more in order to meet their nutritional requirements. Similarly, when nutrient dilution occurs, the number of specialized mining and galling damage types is unlikely to increase, as is the number of occurrences of these damage types. Gall-inducing insects hijack their host plant’s metabolism; their control over the tissues that form galls may well be sufficient to shield them from the effects of nutrient dilution. However, the amount of surface area lost to mining may increase if larvae need to make longer mines in order to fulfill their nutritional requirements. External foliage feeding is the only broad category of herbivory for which one would expect to see the number of feeding event occurrences increase with nutrient dilution. External foliage feeders, like leaf miners, will need to consume more leaf surface area if nutrient dilution reduces the nutritional quality of the leaf blade. And unlike leaf miners, external foliage feeders have the opportunity to partition the additional leaf surface area they consume among multiple feeding event occurrences.

Whereas “damage type diversity” as discussed here is a measure of the richness of damage types, the evenness of herbivore damage also holds potential to inform differences in herbivory among assemblages. Gunkel and Wappler (2015) extended the insights of Olszewski (2004) regarding the relationship between rarefaction and evenness to the study of insect herbivory. A number of other authors have also quantified evenness—of the floral community, of damage types, or both—when evaluating insect herbivory in the fossil record (Wappler et al., 2015; Currano et al., 2019; Azevedo Schmidt et al., 2019). These too can be used as additional dimensions of the herbivory ecospace. Of note, the frequencies of each plant host and damage type can be replaced with their respective surface areas when calculating evenness metrics.

Another metric, related to evenness, is the offset between the prevalence of each plant host and the prevalence of insect damage on it. For example, if a locality contains two plant hosts, one that accounts for 70% of total leaf surface area and another that accounts for 30%, a null expectation would be that 70% of herbivorized surface area would belong to the first plant host and 30% would belong to the second. The extent to which this null expectation is violated can be quantified by summing the absolute values of the differences between each plant host’s prevalence in the assemblage and the proportion of herbivorized leaf area that it contains. In this example, if the first host plant contains 65% of herbivorized leaf area and the second contains 35%, the offset between total and herbivorized leaf area summed across the locality is |0.7 *−* 0.65| + |0.3 *−* 0.35| = 0.1. If, on the other hand, the first host plant contains only 25% of herbivorized leaf area and the second contains 75%, the offset between total and herbivorized leaf area summed across the locality is |0.7 *−* 0.25| + |0.3 *−* 0.75| = 0.9. This evenness-offset score ranges from 0 (no offset) to 2 (maximum offset). This metric can be leveraged as an additional dimension for an herbivory ecospace. An 84% confidence interval for this evenness-offset score can be generated by resampling plants and damage type occurrences, with replacement, to the amount of surface area observed in the original dataset, calculating an evenness-offset score for the resampled datasets, and iterating this process 5,000 times. (For discussion of the number of iterations needed, see the supplemental material.)

Floral diversity, and the relationship of floral diversity to damage type diversity, can comprise additional dimensions of the herbivory ecospace. A damage-diversity-to-floral-diversity ratio, in which rarefied damage type diversity is divided by rarefied floral diversity, holds the potential to reveal the extent to which floral diversity underlies damage type diversity. A confidence interval for this ratio can be calculated by randomly selecting a value within the confidence interval for rarefied damage type diversity, randomly selecting a value within the confidence interval for rarefied floral diversity, dividing the former by the latter, and iterating this process 1,000 times. The number of specimens required to rarefy floral diversity to sample coverage of 0.8 may differ from the number of specimens required to rarefy damage type diversity to sample coverage of 0.8.

Lastly, the prevalences of each functional feeding group within an assemblage can comprise yet more dimensions of the ecospace. Ideally, the prevalence of each functional feeding group would be quantified by the absolute and relative amounts of herbivorized surface area that it represents. However, as noted above, surface area measurements are unavailable for nearly all herbivory datasets. Thus, the proportion of damage type occurrences attributable to each functional feeding group can instead be used as a proxy. Confidence intervals for the proportions of either herbivorized area or damage type occurrences can be calculated by iteratively resampling with replacement as discussed above.

Additional measures, such as bipartite network metrics (*sensu* Blüthgen et al., 2008) and host specificity, require so much data that it may not be possible to precisely and accurately estimate them with a typical fossil herbivory dataset. This is further discussed in the supplemental material (Figure S6; “evaluation of other metrics”).

### 2.4 The optimal number of dimensions in the ecospace

When traditional multivariate statistics are used in ecological studies, scree plots allow researchers to determine the number of axes that warrant interpretation (McGarigal et al., 2013). We are not aware of any analysis comparable to a scree plot that would facilitate determination of whether all of the above dimensions of an ecospace warrant consideration. However, we also do not see any *a priori* reason to expect that all of these dimensions will be informative in all studies of insect herbivory. We have listed all of these dimensions because they are already used in studies of insect herbivory and we aim to standardize statistical practices and to encourage use of the most robust metrics. It is entirely reasonable to assume at the beginning of any study that damage type diversity and the herbivory index are the only necessary dimensions of this ecospace unless another dimension can directly address any hypotheses under consideration. If two or more dimensions of an ecospace are highly colinear, or if one or more dimensions contribute only statistical noise, the most appropriate course of action is to eschew those dimensions.

## 3 The amount of sampling required

The ability to determine *a priori* how much sampling is needed for a given assemblage—or whether an assemblage contains sufficient material for robust comparisons with other assemblages—would allow investigators to allocate their efforts most efficiently. Unfortunately, our results show that neither the number of leaves nor the amount of surface area in an assemblage are reliable predictors of sufficient sampling for coverage-based rarefaction. The frequency of damage on leaves is a far more reliable predictor, but is much more difficult to estimate prior to data collection.

One advantage of coverage-based rarefaction is that it can be integrated with a “stopping rule”. In the case of the sample coverage threshold of 0.8 used here, sample coverage can be re-calculated whenever new data are added to a dataset and sampling can be considered sufficient when the threshold of 0.8 has been reached. Although stoppage of data collection at this threshold would allow a maximum number of assemblages to be examined and compared, it is worth noting that the confidence intervals surrounding damage type diversity and the herbivory index will narrow if sampling continues beyond a sample coverage of 0.8.

As discussed in a recent contribution, estimates of damage type diversity require more sampling than estimates of the herbivory index do (Schachat et al., 2020). As shown in the supplemental material (“evaluation of other metrics”; Figure S6), even more sampling would be needed to estimate host-specificity of individual damage types and to estimate more complex metrics that match particular damage types with particular host plants; the extreme sensitivity of such metrics to sample size has already been established in the neontological literature (Blüthgen, 2010).

In fact, it has yet to be demonstrated that any realistic amount of sampling completeness in studies of fossil insect herbivory is adequate for reliable calculation of complex metrics such as bipartite network properties. The sampling completeness necessary for reliably calculating bipartite network properties could be established by iteratively subsampling an assemblage to half of its original leaf surface area and calculating network properties for each subsample. The results of this subsampling routine could be used to generate a confidence interval for each network property. If, for each network property, the confidence interval is narrow enough that it does not overlap with the confidence intervals generated for other assemblages subsampled to the same amount of surface area, the amount of surface area available is sufficient to reliably calculate network properties. Of note, however, the requisite amount of surface area will vary with the dominance– diversity structure of the plant host and damage type communities at each assemblage. It is not yet clear whether it is worth pursuing sufficient sampling for the implementation of bipartite network analyses.

## 4 Generation of null hypotheses

The study of insect herbivory in the fossil record has become vastly more popular over the past few decades. Early studies that examined patterns of insect herbivory caused by a major event, such as a mass extinction, had only one available option: to quantify the magnitude of the differences in insect herbivory before and after the event in question. A question that could not be addressed at the time those early studies were conducted, but can be addressed now, is whether the amount of change in insect herbivory after a major event exceeds the amount of change that one would expect if the event had not happened.

Returning to the end-Triassic example mentioned in the Introduction, a number of factors preclude the attribution of changes in herbivory across a boundary to the events that occurred at the boundary. First, the difficulty of finding floral assemblages deposited immediately before or after the Triassic/Jurassic boundary, and establishing their proximity to the boundary, necessitates that this event be examined with Triassic and Jurassic floras that are separated by millions of years—a long enough time for plant and insect communities to change without any extraordinary abiotic events. Second, even within a single basin, plant communities, soils, precipitation, temperature, and perhaps also predators and parasitoids change with major abiotic events, introducing various potential causes of changes in insect herbivory that may not be directly related to the abiotic event in question (Currano et al., 2019). On a related note, although nutrient dilution is expected to cause an increase in the herbivory index, several of these alternative phenomena could also cause such an increase. Third, extant ecosystems demonstrate that coeval plant communities in close geographic proximity can vary tremendously due to microclimate and other factors such as soil type and frequency of disturbance (Tamme et al., 2010); the difficulty of identifying a characteristic flora for a particular place and time makes it more difficult still to compare characteristic patterns of insect herbivory for a particular place across different time slices.

To overcome these obstacles, many studies of insect herbivory have examined multiple plant assemblages from each time interval of interest (Labandeira et al., 2002a; Currano et al., 2008; Wappler, 2010; Donovan et al., 2018). If the assemblages from each time interval form distinct clusters, in an NMDS plot for example, this provides far stronger evidence of change over time than can be gleaned from only two assemblages. The outstanding question is how much change over time is to be expected in the absence of an event such as an environmental disturbance (the amount of change expected as a null hypothesis), and how much change can be attributed to the event under consideration (the amount of change needed to support the alternative hypothesis that the event in question caused significantly more change in patterns of insect herbivory than can be explained only by the passage of time).

The theoretical ecospace outlined here holds the potential to go beyond an examination of the number of damage types observed per number of leaves, by quantifying the following. First is the amount of variation in insect herbivory to be expected among coeval floras that occur in close proximity and have slightly different plants, soils, depositional conditions, and microclimates. Second is the amount of variation in insect herbivory to be expected on the time scales examined in relevant studies (10 Kyr, 100 Kyr, 1 Myr, 10 Myr) in the absence of any major environmental changes. Third is the amount of, and the nature of, changes in insect herbivory associated with environmental changes, such as the Cretaceous/Paleogene Event (Labandeira et al., 2002b; Donovan et al., 2018) and the Paleocene/Eocene Thermal Maximum (Currano et al., 2008). Fourth is the amount of, and the nature of, changes in insect herbivory associated with insect evolution over tens of millions of years. Nearly any comparison of floral assemblages will yield evidence of different patterns of insect herbivory; the key question is how much change needs to occur in order to rise above background levels. In the future, as data become available to include increasing numbers of assemblages in this theoretical ecospace, confidence intervals can be established in each dimension for these four types of changes in insect herbivory.

Traditional morphospaces have already been leveraged to generate null hypotheses and to disentangle the results of directional evolution from changes through time that are best attributed to a “random walk” model (Pie and Weitz, 2005; Puttick et al., 2020). The same logic can be extended to the study of insect herbivory in the fossil record.

## 5 Preliminary results

### 5.1 Comparable herbivory during the paleozoic, mesozoic, and cenozoic

When damage type diversity from the assemblages in Tables 1, 2, and 3 is plotted against time, the most striking result is the overlap of damage type diversity among Permian, Triassic, and angiosperm-dominated assemblages (Figure 5). The dataset analyzed here contains five times as many angiosperm-dominated assemblages as it does Paleozoic assemblages, and over three times as many angiosperm-dominated assemblages as Triassic assemblages. And yet the Laguna Polina assemblage from the Permian of Patagonia (Cariglino, 2018), the Kühwiesenkopf assemblage from the Triassic of Italy (Labandeira et al., 2016), and Cyphergat 111A assemblage from the Triassic of South Africa (Labandeira et al., 2018) all contain damage type diversities within the highest 12% of values for angiosperm-dominated assemblages. The supplemental material includes a sensitivity analysis and a discussion of the role of rare damage types in diversity estimates.

**Figure 5:**
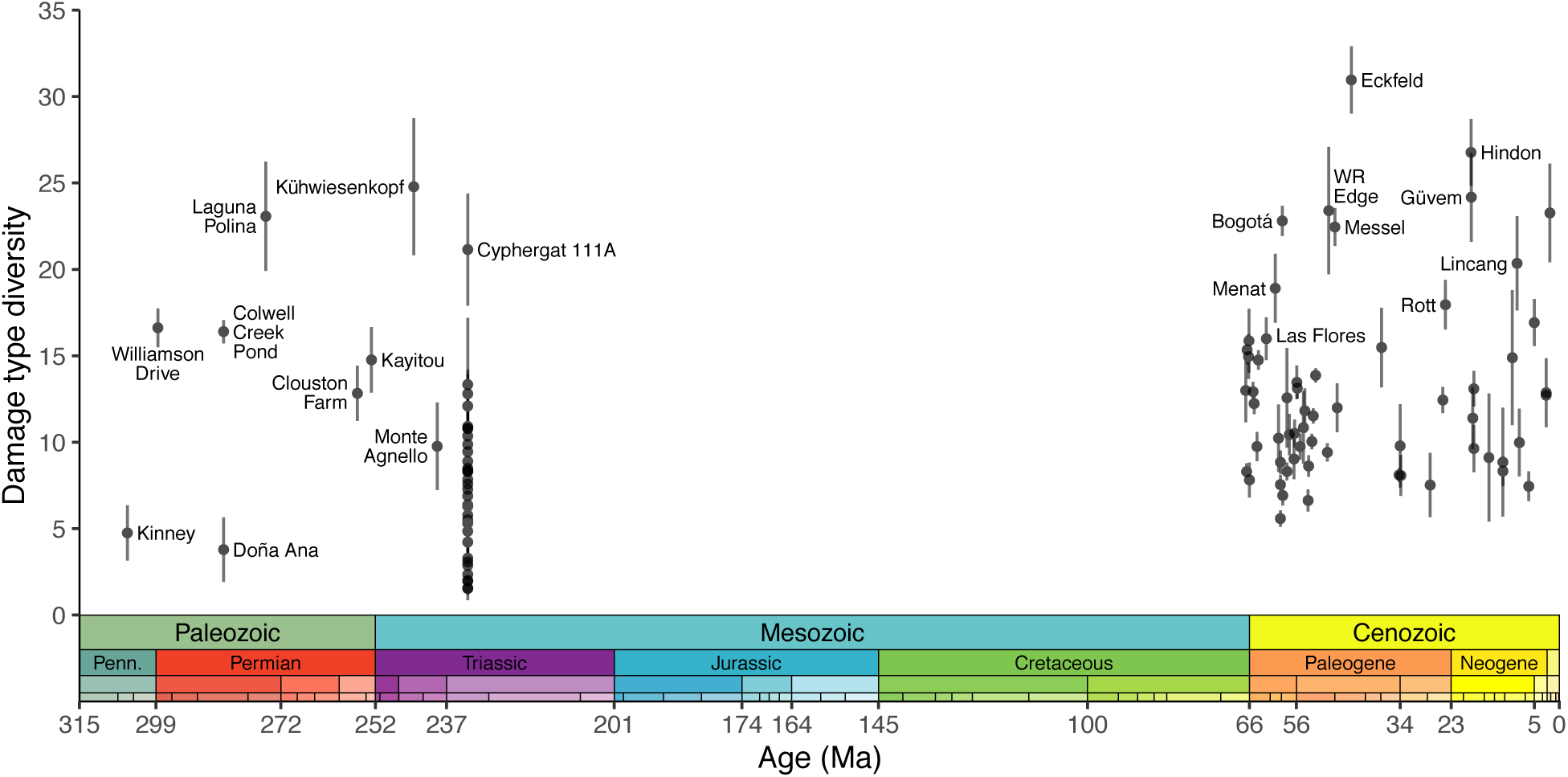
Damage type diversity estimated with coverage-based rarefaction, plotted against time.

The minimal increase in the upper bound of damage type diversity from the Paleozoic through Cenozoic is further supported by the two-dimensional ecospace which we advocate for here (Figure 6). The available data are sufficient to include only eight assemblages in this ecospace: Williamson Drive (Xu et al., 2018) and Colwell Creek Pond (Schachat et al., 2015) from the late Paleozoic; and Daiye Spa, Hubble Bubble, PN, Fifteenmile Creek, Republic, and Bonanza from the interval surrounding the Paleocene/Eocene Thermal Maximum (Wilf et al., 2001; Labandeira, 2002; Currano et al., 2008, 2010, 2016). This ecospace also supports the prediction outlined earlier in this contribution that an abiotic event—in this case, exemplified by the Hubble Bubble locality during the PETM—can indeed lead to a change in damage type diversity, but most clearly manifests with a change in the herbivory index.

**Figure 6:**
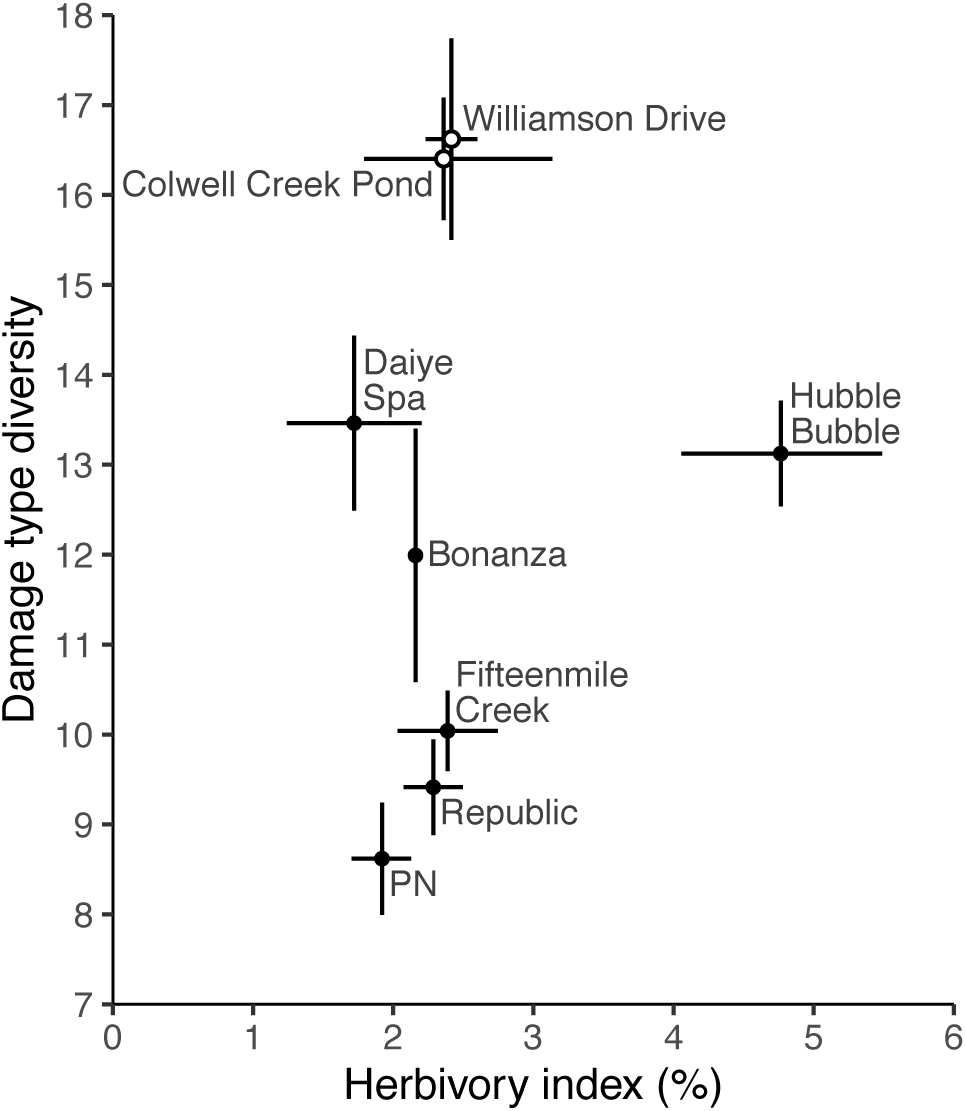
Damage type diversity plotted against the herbivory index in a two-dimensional ecospace. Paleozoic localities are denoted with white circles and angiosperm localities are denoted with dark circles. The confidence intervals for the herbivory index were derived with the method of Schachat et al. (2018) for Williamson Drive and Colwell Creek Pond, and with the method of Currano et al. (2016) for all other assemblages.

This finding of similar damage type diversities in Paleozoic, Triassic, and angiosperm-dominated assemblages may be somewhat surprising given the angiosperm potential for uniquely high productivity (Boyce and Zwieniecki, 2012) and their tremendous diversity (Soltis and Soltis, 2004)—and especially their codiversification with insects (Pellmyr, 1992). Leaf mining, for example, is a highly specialized form of herbivory that accounts for a wide array of described damage types (Eiseman, 2019) and occurs overwhelmingly on angiosperm leaves (Scott et al., 1992). One might expect the origin of angiosperms to have caused a diversification of leaf-mining insects, which in turn would have caused an increase in damage type diversity. However, the data do not support an overall increase in damage type diversity in angiosperm-dominated assemblages.

The difficulty of quantifying the role of angiosperm diversity and biomass in shaping extant ecosystems stems largely from their overwhelming dominance. If angiosperms accounted for 70% of the diversity and biomass of land plants, any disproportionate ecological role they may fulfill—for example, if they hosted 95% of leaf mine damage types—would be easier to establish. But, because angiosperms account for approximately 96% of described vascular plant species (Christenhusz and Byng, 2016), any disproportionate ecological role they may fulfill—for example, if they hosted 98% of leaf mine damage types—may be statistically indistinguishable from the null expectation of this ecological role (in the case of this example, that angiosperms would host 96% of leaf mine damage types). To use another hypothetical example, if it were known with absolute certainty that angiosperms host only 93% of leaf mine damage types, one could still say that the overwhelming majority of leaf mine damage types occur on angiosperms, such that leaf mine diversity and thus overall damage type diversity likely increased alongside the radiation of flowering plants.

### 5.2 The role of *p*CO_2_ depends on the rate of atmospheric change

The changes in damage type diversity in Figure 5 show no general relationship with *p*CO_2_. For example, the Kayitou assemblage dates to the late Lopingian, an interval for which many geochemical models reconstruct high *p*CO_2_ (Mills et al., 2019), but Kayitou does not have particularly prevalent insect damage (Liu et al., 2020) nor is its damage type diversity unusually high (Figure 5).

Rising *p*CO_2_ may drive nutrient dilution and, thus, increased insect herbivory, but stable *p*CO_2_—whether high or low—may favor instead compensatory physiological changes that allow for stable rates of carbon fixation under different atmospheres. The carbon cycle perturbation of the PETM may have constituted an atmospheric change too rapid for plants to adapt to; it is under this circumstance that nutrient dilution may be most likely to occur. For this reason, the increased intensity of herbivory seen at the PETM is not a template for the changes in herbivory that would occur if *p*CO_2_ rose slowly enough for plants to remain well adapted to the atmospheric composition.

## 6 Considerations for future studies

Overlapping damage type diversities through time are far easier to demonstrate than changing diversities. Labandeira et al. (2002b) evaluated damage type diversity before and after the end-Cretaceous event. Their study included 17 beds with 200 or more dicot leaves: eleven Cretaceous and six Paleogene. Coverage-based rarefaction supports the authors’ conclusion that damage type diversity was higher during the latest Cretaceous than during the earliest Paleogene. However, damage type diversities vary widely within each of these two intervals. These fossil beds all occur within the same basin, and the beds from each interval are dominated by a single type of depositional environment. Aside from the first-order finding of less herbivory in Paleogene beds, variation in damage type diversity does not occur in a predictable manner through time.

This variability observed within 1 Myr in a single basin highlights the difficulty of establishing a representative amount of damage type diversity for a particular interval—even for a single depositional environment within a single basin during a very small amount of geologic time. The low signal-to-noise ratio among the damage type diversities for a pooled set of assemblages highlights the difficulty of identifying temporal trends in insect herbivory.

Additional factors further conceal temporal trends. First is the obvious variability among assemblages in geographic location, insect taxa, plant taxa, soil type, and climate. Second is uneven sampling across time. No studies of herbivory for an entire flora have been published for the Jurassic, and within the Cenozoic, the Paleogene is far better sampled than the Neogene or Quaternary. Third, especially for the Cenozoic, assemblages are chosen for study based largely on their relevance to events such as the end-Cretaceous extinction (Labandeira et al., 2002b; Wilf et al., 2006; Wappler et al., 2009; Donovan et al., 2017, 2018), the Paleocene/Eocene Thermal Maximum (Wilf and Labandeira, 1999; Wilf et al., 2001; Currano et al., 2008; Currano, 2009; Currano et al., 2010), and the Early Eocene Climatic Optimum (Currano et al., 2016, 2019). The assemblages for which we have data are disproportionately likely to come from the least representative part of a geologic interval, such as the very earliest Eocene, because these time slices epitomize the phenomena that underlie paleontologically interesting ecological and evolutionary questions. The uneven temporal distribution of studied assemblages adds far more variability to reconstructed long-term trends than one would expect if assemblages were chosen for study in an unbiased manner, and has the potential to obscure these trends.

## 7 Conclusions

As the proliferation of free software packages and online databases continues, it becomes easier to run analyses and generate graphs with insect herbivory data—regardless of whether the methods are appropriate for the data and regardless of whether the datasets are sufficiently complete to meet the assumptions of the methods. As shown here, the Chao1 estimator, a method that is particularly well-suited to handle the sparseness of insect herbivory datasets, rarely provides estimates of damage type diversity that are both precise enough to minimize the frequency of false negative results and accurate enough to contain the true, asymptotic value.

Moreover, asymptotic damage type diversity calculated through a combination of rarefaction and the Chao1 estimator is formulated as an unbiased estimator but is severely biased by sample size, presumably due to the exceptional sparsity of herbivory datasets. This finding highlights the need to regress estimators of insect herbivory against the number of leaves and damage types in raw datasets to verify that they are not biased by sampling effort.

Whereas size-based rarefaction and the Chao1 estimator present insurmountable limitations for insect herbivory data, coverage-based rarefaction holds the most promise for generating estimates of damage type diversity that are unbiased, and robust to leaf size and sample size. Combined with the herbivory index, which measures the intensity of insect herbivory, estimates of damage type diversity can be leveraged as the foundation of a theoretical ecospace. This ecospace holds the potential to address two overarching questions that remained unresolved for two decades. First is how to distinguish the minor variation in insect herbivory that inevitably occurs among assemblages from slightly different times or places (null hypothesized amount of variation) from the changes in insect herbivory that accompany major abiotic events or innovations in insect evolution (alternative hypothesis). Second is how to distinguish among abiotic causes (e.g., mass extinctions) and biotic causes (insect and plant evolution) of major changes in the amount of insect herbivory.

There are two ways to build upon the many fossil herbivory datasets amassed over the past few decades: collecting more data, and interrogating existing data to develop and refine analytical frameworks. By advancing the latter, a theoretical ecospace for insect herbivory will hopefully underscore the importance the former.

## 8 Acknowledgements

This contribution was made possible by the many paleontologists who have collected fossil plants and described the insect damage they contain, with John Anderson and Heidi Holmes particularly deserving of recognition here. Bárbara Cariglino, Mónica R. Carvalho, Michael P. Donovan, L. Alejandro Giraldo, Christian Müller, Manuel Robledo, Torsten Wappler, and Peter Wilf provided access to datasets. The Paleobotany Reading Group at Stanford University provided valuable discussion. We thank two anonymous reviewers for their very helpful feedback. S.R.S. received funding from the Coleman F. Fung Interdisciplinary Graduate Fellowship, Vice Provost of Graduate Education, Stanford University; from the Harriet Benson Fellowship Award, Department of Geological Sciences, Stanford University; and from the Winifred Goldring Award, Association of Women Geoscientists and Paleontological Society.

## 9 Supplemental material

### 9.1 Difficulties in estimating damage type diversity

The differences in sampling intensity among Colwell Creek Pond and Mitchell Creek Flats pose an insurmountable obstacle to estimating true damage type diversity with raw or rarefied data. Rarefaction curves that include the dominant broadleaf plant taxa from both assemblages show that the estimate of higher damage type diversity for Colwell Creek Pond cannot be disentangled from the disparities in the amount of surface area examined (Figure S1A). If the damage type diversities tallied at each assemblage, 45 at Colwell Creek Pond and 19 at Mitchell Creek Flats, are interpreted at face value, the resulting conclusion that former contains greater damage type diversity is based entirely on an artifact of uneven sampling. However, even when the role of sampling completeness is taken into account, the overlapping confidence intervals complicate estimates of whether greater sampling would reveal significant differences in damage type diversity among the two assemblages (Knezevic, 2008).

The unreliability of diversity estimates derived from incompletely sampled assemblages is further underscored by the variability of estimated damage type diversity at Colwell Creek Pond when rarefaction curves are calculated with leaves from this locality that are randomly sampled only to the amount of surface area seen at Mitchell Creek Flats (Figure S1B–D). In other words, the difficulty of extrapolating damage type diversity at Mitchell Creek Flats for the amount of leaf area sampled from Colwell Creek Pond is due not only to the uncertainty surrounding damage type diversity beyond the amount of leaf area available, but also to the uncertainty surrounding the reliability of the damage type diversity estimated for Mitchell Creek Flats at the amount of leaf area that has been examined for this assemblage. Comparisons of damage type diversity, therefore, require a method such as rarefaction to control for differences in sampling completeness, and require fairly complete sampling of all localities—which can be difficult to achieve due to both the availability of fossil material and the investigator effort required.

The difficulty of estimating damage type diversity in light of sampling incompleteness is further compounded by the unavailability of surface area data for nearly all assemblages examined thus far (Schachat et al., 2018). Because leaf size varies so widely, two assemblages can have equivalent numbers of damage types per 400 cm^2^ of leaf surface area while having widely different numbers of damage types per 400 leaves. Variable leaf surface area can complicate, if not invalidate, attempts to use damage type data to discern macroevolutionary and macroecological patterns. For example, the amount of surface area per leaf available to insect herbivores has changed through time because gymnosperms and angiosperms have different average leaf sizes. As another example, whereas heightened levels of insect herbivory in the tropics form the basis of various biogeographical theories (Dobzhansky, 1950; MacArthur, 1969; Janzen, 1970; Connell, 1971), the magnitude of the latitudinal variability in herbivory is not entirely understood and depends considerably on the metric used (Anstett et al., 2014; Andrew and Hughes, 2005; Adams et al., 2011, 2009; Zhang et al., 2011; Adams and Zhang, 2009; Adams et al., 2010; Salazar and Marquis, 2012; Moreira et al., 2015; Moles et al., 2011). And because plants in wet biomes tend to have larger leaves (Ackerly et al., 2002; Cunningham et al., 1999; Givnish, 1987), rarefaction curves that are scaled by number of leaves rather than amount of leaf surface area may confound increased damage type diversity in wet tropical forests with increased leaf size in these habitats.

In addition to subsampling techniques such as rarefaction, a number of population estimators hold the potential to calculate true (asymptotic) damage type diversity and corresponding confidence intervals from datasets of varying levels of completeness (Chao, 1987; Chiu et al., 2014; O’hara, 2005; Palmer, 1990; Smith and van Belle, 1984). Of these, the Chao1 estimator is particularly well-suited to sparse datasets (Chao, 1989). However, these estimators follow the same pattern seen in rarefaction curves of steeply increasing estimates of diversity at low levels of sampling completeness (Chao et al., 2009). One might hope that the Chao1 estimator would behave like estimates of the herbivory index, with confidence intervals that contain the true value even when sampling is very incomplete and then narrowing more closely around the true value as sampling becomes more complete. However, this often is not the case. An analysis of plant assemblages for which at least 1,000 broadleaf specimens have been examined for insect herbivory (Tables 1, 2) shows that confidence intervals are deceptively narrow when sampling is insufficiently complete to capture the true value of damage type diversity (Figure S2). As sampling increases, the confidence intervals are more likely to contain an accurate result but often widen so much that false negative results become inevitable (Figure S2).

Whereas neither rarefaction nor the Chao1 estimator provide sufficient accuracy and precision to estimate damage type diversity, they have the potential to work far better in concert. The combination of rarefaction and Hill numbers (Chao et al., 2014), a family of metrics to which the Chao1 estimator belongs, was an extension of advances in the extrapolation of rarefaction curves (Colwell et al., 2012) and in rarefaction based on sampling completeness rather than sample size (Chao and Jost, 2012). The Chao1 estimator return an identical point estimates and confidence intervals for asymptotic diversity for a given frequency of damage type occurrences regardless of whether those damage type occurrences are spread out over 10 or 100,000 leaves. But, unlike the Chao1 estimator, rarefaction has the advantage of taking sampling completeness into account while generating diversity estimates. Unlike rarefaction, the Chao1 estimator has the advantage of robustness to differences in leaf size and fragmentation because this metric estimates true, asymptotic diversity rather than diversity at a particular level of incomplete sampling. The method of Chao et al. (2014) possesses both of these advantages and is implemented in the R package iNEXT (Hsieh et al., 2016).

However, the method of Chao et al. (2014) yields estimates of asymptotic damage type diversity that are biased by the number of leaves sampled. This is probably because of the extreme sparsity of damage type occurrence datasets combined with the large quantity of damage types that are only observed once. The sensitivity of this method to sampling completeness can be seen when comparing the rarefaction curves for assemblages with over 7,000 leaves examined to extrapolated rarefaction curves generated after subsampling each assemblage down to 1,000 or 2,000 leaves (Figure S3). 2,000 leaves are rarely sufficient to extrapolate the complete rarefaction curve, and 1,000 leaves are sufficient for only one assemblage, Willershausen (Adroit et al., 2018). With 1,000 leaves, these extrapolated rarefaction curves level off far too quickly—and even for Willershausen, the one assemblage that does not follow this pattern, the 95% confidence interval is much too wide to permit the detection of significant differences in damage type diversity among most assemblages.

### 9.2 The unreliability of extrapolated diversity estimates generated with coverage-based rarefaction

To evaluate whether rarefied damage type diversity at a sample coverage of 0.8 can be extrapolated from datasets that do not reach this level of coverage, we used an iterative subsampling procedure. Leaves were subsampled from the pre-angiosperm assemblage with the largest number of leaves examined, Aasvoëlberg 411 (Labandeira et al., 2018), and the angiosperm assemblage with the largest number of leaves examined, Willershausen (Adroit et al., 2018). This procedure was repeated 3,000 times for each assemblage to ensure that a wide range of levels of sample coverage are represented among the subsampled datasets. For each subsampled dataset, we extrapolated rarefied damage type diversity to a sample coverage of 0.8. The results of this procedure (Figure S4) show that the accuracy of extrapolated estimates of damage type diversity does increase with sample coverage, but these extrapolated estimates are never accurate enough to inspire any confidence. Therefore, we do not recommend extrapolating estimates of damage type diversity for assemblages with a sample coverage below 0.79.

### 9.3 Evaluation of other metrics

The ecological literature contains a variety of complex metrics for the evaluation of trophic interactions. With paleontological data, and especially paleontological data from a clade with a history of biased collecting (Gunkel and Wappler, 2015), a fundamental question is whether the data at hand are sufficient to yield robust results when analyzed with more complex techniques.

Both plants and their damage types follow a dominance-diversity distribution that approximates a lognormal or gamma distribution (Figure S5), as do nearly all other biotic communities (Diserud and Engen, 2000). In a typical floral assemblage, the majority of plant hosts and damage types are rare.

Generating a standardized estimate of floral or damage type diversity at a given assemblage through rarefaction is a relatively simple matter of subsampling the more abundant taxa in a consistent fashion. Estimating relationships among plant hosts and damage types, however—for example, estimating the proportion of damage types on the second-most abundant plant host which also occur on the most abundant plant host—is a far more complex matter, and thus is far more sensitive to sample size. A single tree can grow thousands of leaves in a single year (White, 1993), but it is uncommon for thousands of leaves from a single fossil assemblage to be examined for insect herbivory (Tables 1 and 2). We are not sure if all fossil leaf material ever examined for the herbivory index contains more or less leaf surface area than a single elm tree.

Therefore, whereas the amount of sampling typically seen in studies of fossil herbivory is likely sufficient to determine whether the most common damage type at an assemblage occurs on the most common plant host at the assemblage, this amount of sampling is probably not sufficient to determine whether the twentieth-most-common damage type occurs on the twentieth-most-common plant host. The absence of the twentieth-most-common damage type on the twentieth-most-common plant host may indicate that this particular interaction did not occur in the community represented by the fossil assemblage, but can just as easily indicate that sampling is not sufficient to document this interaction.

To evaluate the reliability of complex metrics of insect herbivory calculated for fossil assemblages, we conducted two iterative subsampling routines with data from the pre-angiosperm assemblage with the largest number of leaves examined, Aasvoëlberg 411 (Labandeira et al., 2018), and the angiosperm assemblage with the largest number of leaves examined, Willershausen (Adroit et al., 2018). In both routines, we subsampled 1,000 leaves per iteration and iterated this procedure 1,000 times. For the first routine, we calculated the proportion of damage types occurring on the second-most-abundant plant host that also occur on the most abundant plant host. (This is perhaps the simplest metric of the nestedness of damage type communities among the plant hosts at an assemblage.) For the second routine, we divided the number of leaves belonging to the second-most-abundant plant host by the number of leaves belonging to the most abundant plant host. (The simplicity of this metric is emblematic of the metrics commonly used in the study of fossil herbivory.)

The results of these subsampling routines indicate that, whereas the dominance-diversity structure of the plant hosts within an assemblage is a valid and reliable metric (Figure S6B), the nestedness of damage type communities among individual plant hosts is not (Figure S6A). No sensitivity analyses are needed to demonstrate the impossibility of determining with any certainty whether the twentieth-most-common damage type at an assemblage is truly absent from the twentieth-most-common plant host—as opposed to evading detection due to the number of leaves examined. One could argue that this sort of failure of detection will be shared across all assemblages and therefore will not bias comparisons among assemblages. However, our sensitivity analysis demonstrates that even the very simplest measurement of damage type nestedness (the degree of nestedness among the two most abundant plant hosts) is hopelessly unreliable with 1,000 leaves.

Some analytical trends in paleobiology follow a boom–bust cycle in which a technique increases in popularity, perhaps due to positive connotations associated with its complexity, only to fall out of favor when its reliability, validity, and interpretability come into question (Smith et al., 1997). Our results indicate that the complexity of techniques that associate particular damage types with particular host plants, impressive as they may seem, require far more complete sampling than can be expected from studies of insect herbivory on fossil leaves.

These results also have implications for quantifying host specificity. At present, it is customary to rank each damage type within an assemblage by its host specificity on a discrete scale of generalist (1), intermediate specificity (2), or specialized (3). To avoid misinterpreting artifacts of incomplete sampling as biological phenomena, these rankings are only assigned to damage types that occur on three or more plant specimens. However, the findings presented in this section raise the question of whether three occurrences of a damage type are sufficient to determine its host specificity, particularly if a damage type is classified as specialized because all three of its occurrences are on the most abundant plant host at the assemblage.

### 9.4 Rare damage types and a comparison of angiosperm- and non-angiosperm-dominated assemblages

With size-based rarefaction, one could argue that higher damage type diversities in angiosperm-dominated floras might become apparent at higher sample sizes. An analogous argument for coverage-based rarefaction is that higher damage type diversities in angiosperm-dominated floras might become apparent at higher levels of sample coverage. To evaluate this possibility, we repeated all coverage-based rarefaction analyses by rarefying to a sample coverage of 0.9 instead of 0.8. This higher level of sample coverage incorporates a greater number of rare damage types in diversity estimates, thus constituting a sensitivity analysis of whether the similar damage type diversities estimated for Permian and Cenozoic assemblages in Figure 5 are attributable to the level of sample coverage to which the assemblages were rarefied.

For each assemblage we calculated the difference in damage type diversity when rarefied to a sampling coverage of 0.8 of 0.9 (Figure S7). We found that the differences are smallest for the Molteno assemblages listed in Table 1, which contain relatively low damage type diversities. Although far more angiosperm-dominated assemblages have been evaluated for insect herbivory than non-angiosperm-dominated assemblages outside of the Molteno Formation, the extent to which estimated damage type diversity differs with sample coverage is similar among these two categories. Of note, when sample coverage increases from 0.8 to 0.9, the four assemblages with the greatest increase in estimated damage type diversity are the Hindon Maar, Eckfeld, Messel, and Bogotá assemblages, all of which are angiosperm-dominated and contain high damage type diversities. To discern whether this great increase in damage type diversity with heightened sample coverage is truly unique to angiosperm-dominated assemblages, far more non-angiosperm-dominated assemblages will need to be evaluated for insect herbivory.

Insect herbivory from these four assemblages was described within the last ten years (Wappler et al., 2012; Möller et al., 2017; Giraldo et al., 2021). Estimates of damage type diversity may be biased toward assemblages that have been described most recently under a scenario in which existing damage types are have increased in number (Dos Santos et al., 2020; Xiao et al., 2021). However, Giraldo et al. (2021) found that this is not the case: a more conservative approach toward splitting damage types does not yield a noticeable decrease in damage type diversity as standardized through rarefaction.

### 9.5 The number of iterations needed for resampling procedures

To determine the number of iterations needed for the resampling procedures discussed in “Evaluating other potential dimensions of an ecospace for herbivory,” we iteratively performed our suggested resampling routine for quantifying uncertainty surrounding the offset between the prevalence of each plant host and the prevalence of insect damage on it. For this procedure we used data from the Colwell Creek Pond assemblage (Schachat et al., 2014) because surface area measurements are available. We found that the width of the 84% confidence interval varies minimally whether 100 or 100,000 iterations of the resampling routine are performed, but stabilizes around 5,000 iterations (Figure S8). Therefore, we recommend that future studies use 5,000 resampling iterations to generate confidence intervals.

### 9.6 Criteria for inclusion of leaves and damage types

As noted in the main text, C.C. Labandeira employs a wide definition of “foliage” that includes needles, liverworts, phyllids, photosynthetic wings of seeds, and even flattened horsetail axes whereas S.R. Schachat employs a narrower definition restricted to multi-veined broad leaves and leaves with a defined midvein. (Workers who study angiosperm assemblages typically examine leaves that are at least 50% complete. No specimens were removed from the raw datasets for any of the angiosperm-dominated assemblages analyzed here.) When deciding which specimens from published datasets (Prevec et al., 2009; Cariglino, 2018; Labandeira et al., 2018; Liu et al., 2020; Bernardi et al., 2017) to include in the analyses presented here, we employed a compromise definition that excludes needles, liverworts, phyllids, photosynthetic wings of seeds, and flattened horsetail axes but includes scale leaves.

The raw data for the Williamson Drive assemblage (Xu et al., 2018) preclude determinations of whether many of the individual specimens represent, or at least contain, broadleaf foliage. Therefore, the plant hosts from this assemblage included here are the five foliage types that were included in NMDS plot in the original publication: *Sigillariophyllum* leaves, *Pseudomariopteris cordato-ovata*, *Annularia carinata*, *Lilpopia raciborskii*, and *Macroneuropteris scheuchzeri*.

The data for the Wuda flora (Feng et al., 2020) do not include assignments of damage types. Therefore, damage types were tallied based on the descriptions in the text of the article. An exact count of broadleaf specimens was unavailable due to the vast number of specimens examined. Because of the sparsity of insect damage at Wuda, the amount of sample coverage for this entire dataset is 0.675—well short of the threshold used here, 0.8. Our decisions about the Wuda plant taxa, therefore, are inconsequential, as this assemblage cannot be included in Figure 5.

The description of herbivory at the Lincang assemblage (Zhang et al., 2018) includes a number of unrecognizable damage types. Because the abstract of Zhang et al. (2018) contains a count of damage types that is restricted to those with numbers assigned in the Damage Guide (Labandeira et al., 2007), we followed the authors’ lead and included only those damage types in our analysis.

**Figure S1:**
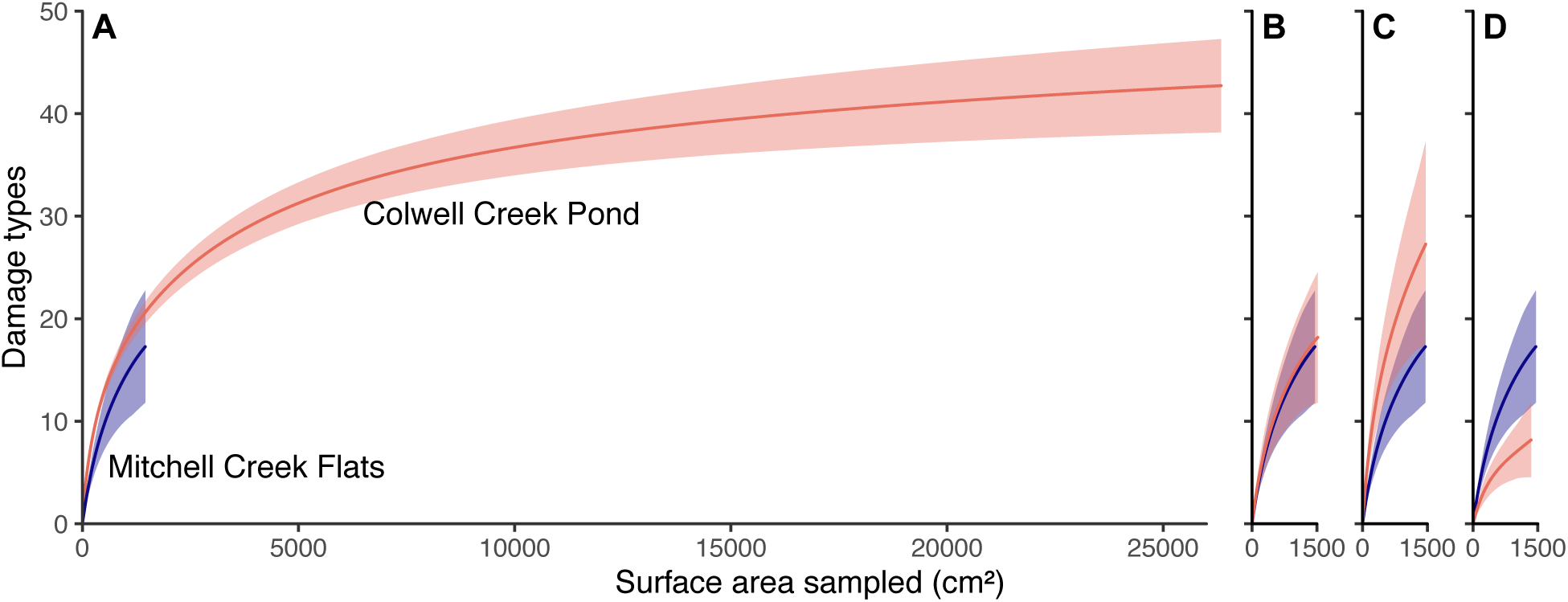
Rarefaction curves for two very similar assemblages: Colwell Creek Pond and Mitchell Creek Flats. **A** shows the rarefaction curves for all data from both assemblages. In **B** through **D**, the rarefaction curves for Colwell Creek Pond are calculated from a randomly subsampled set of leaves with nearly equal surface area to that measured at Mitchell Creek Flats.

**Figure S2:**
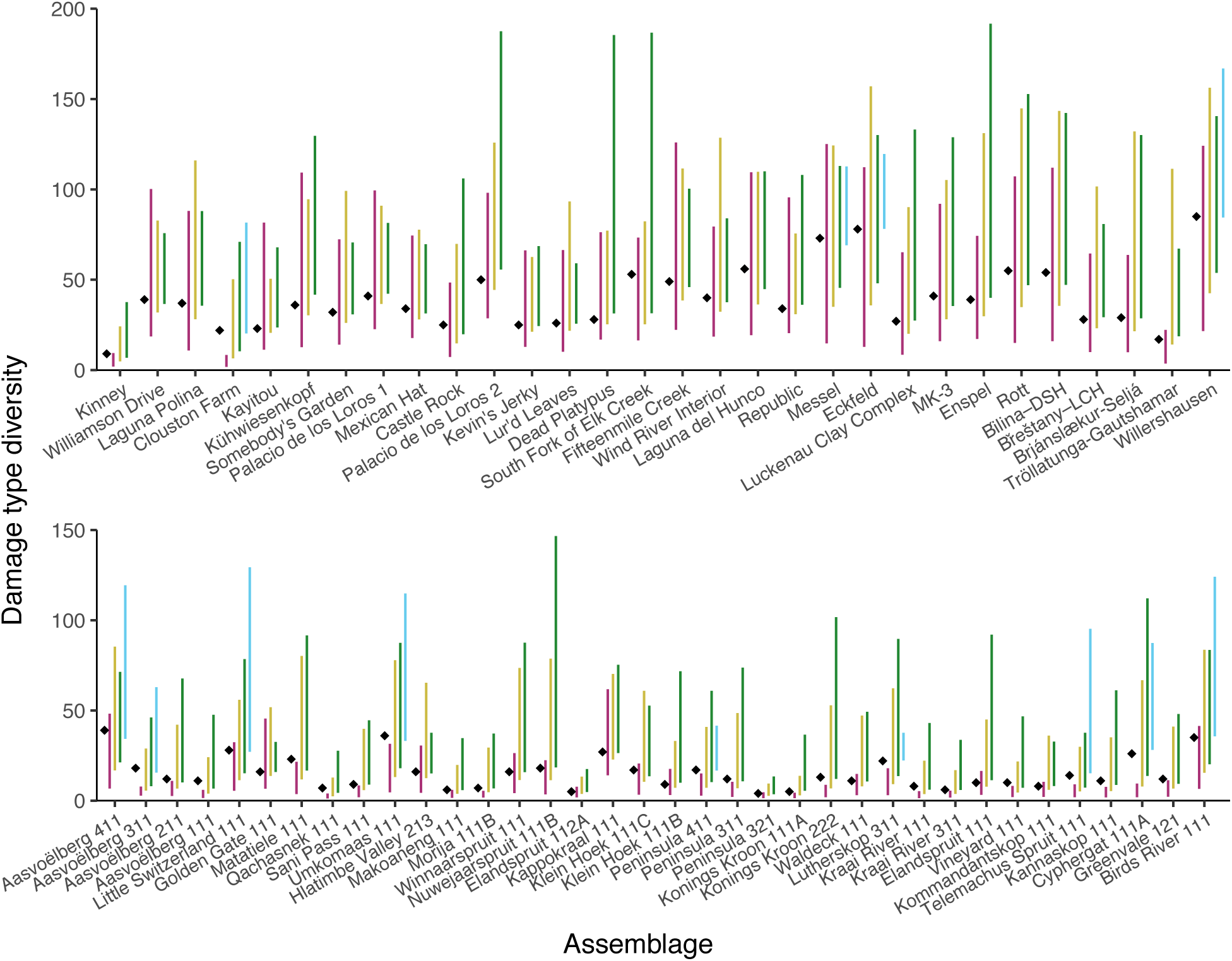
Precision and accuracy of the Chao1 estimator, examined by subsampling all assemblages listed in Tables 1 and 2. The black diamond denotes the raw damage type diversity observed at each assemblage and the lines denote the mean 95% confidence interval of the Chao1 estimator when used on a randomly sampled subset of leaves from the assemblage, as follows: magenta, 100 leaves; yellow, 500 leaves; green, 1,000 leaves; blue, 5,000 leaves. Not all assemblages contain sufficient material to subsample to 5,000 leaves. At low levels of sampling, the Chao1 estimator clearly underestimates damage type diversity. At higher levels of sampling, the Chao1 estimator becomes more accurate but far less precise.

**Figure S3:**
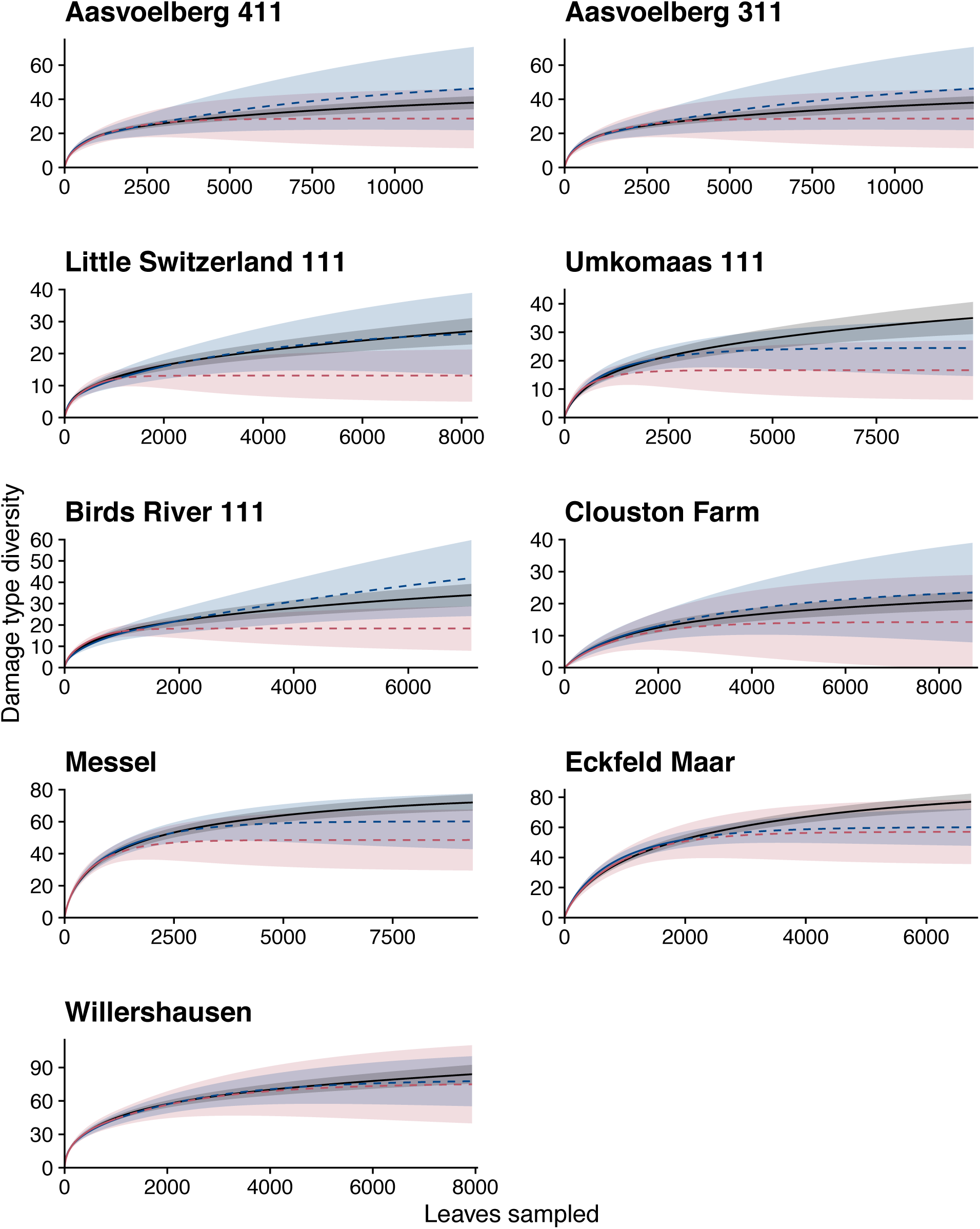
Precision and accuracy of estimates of damage type diversity generated with the method of Chao et al. (2014). Interpolated rarefaction curves are in solid lines and extrapolated curves are in dashed lines. The black lines represent the interpolated rarefaction curve for the raw dataset, the blue lines represent the raw dataset subsampled down to 2,000 leaves and then extrapolated to the number of leaves present in the raw dataset, and the red lines represent the raw dataset subsampled down to 1,000 leaves and then extrapolated to the number of leaves present in the raw dataset.

**Figure S4:**
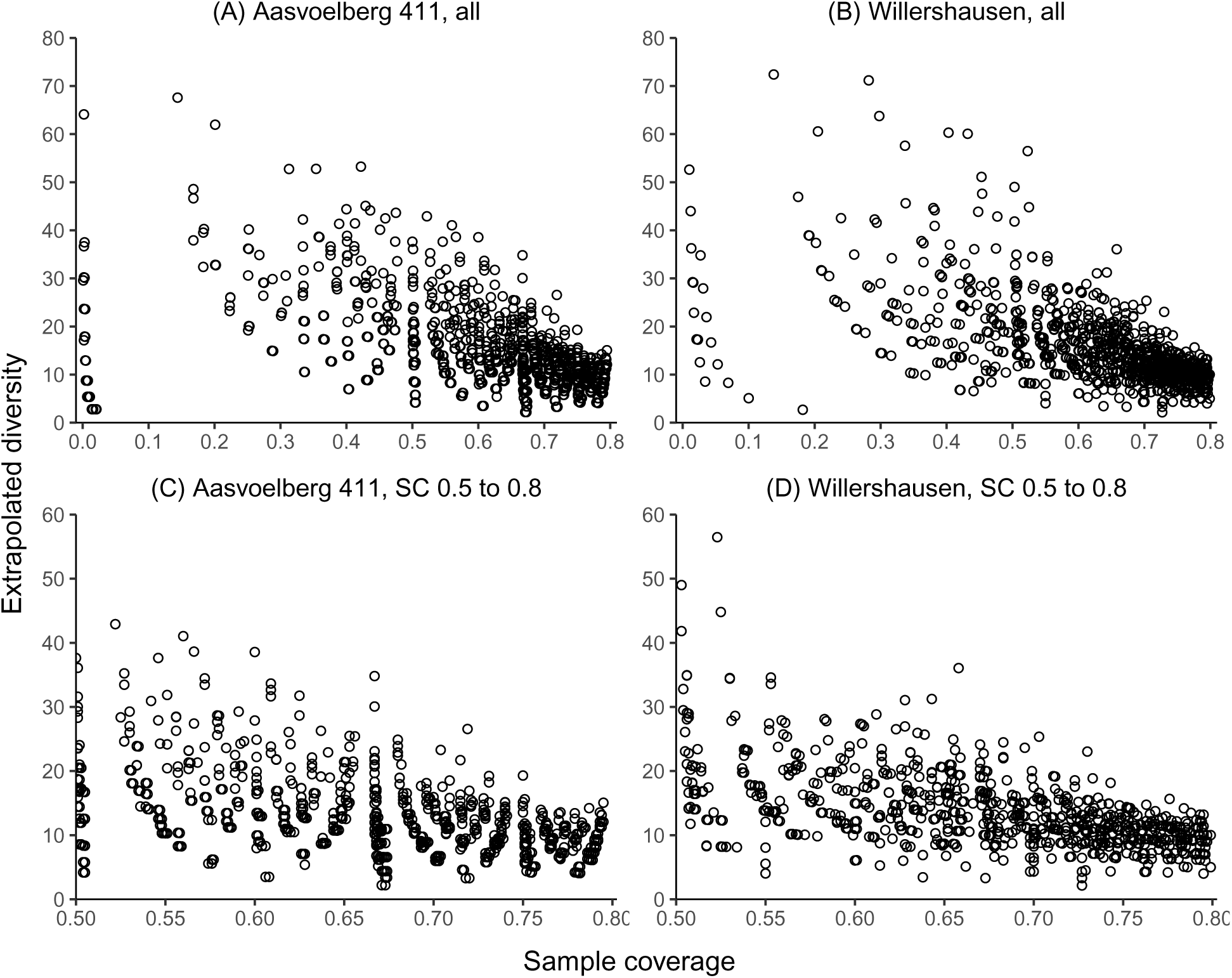
The variability of estimates of damage type diversity at a sample coverage of 0.8 when extrapolated from 1,000 subsampled leaves that yield a sample coverage below 0.8.

**Figure S5:**
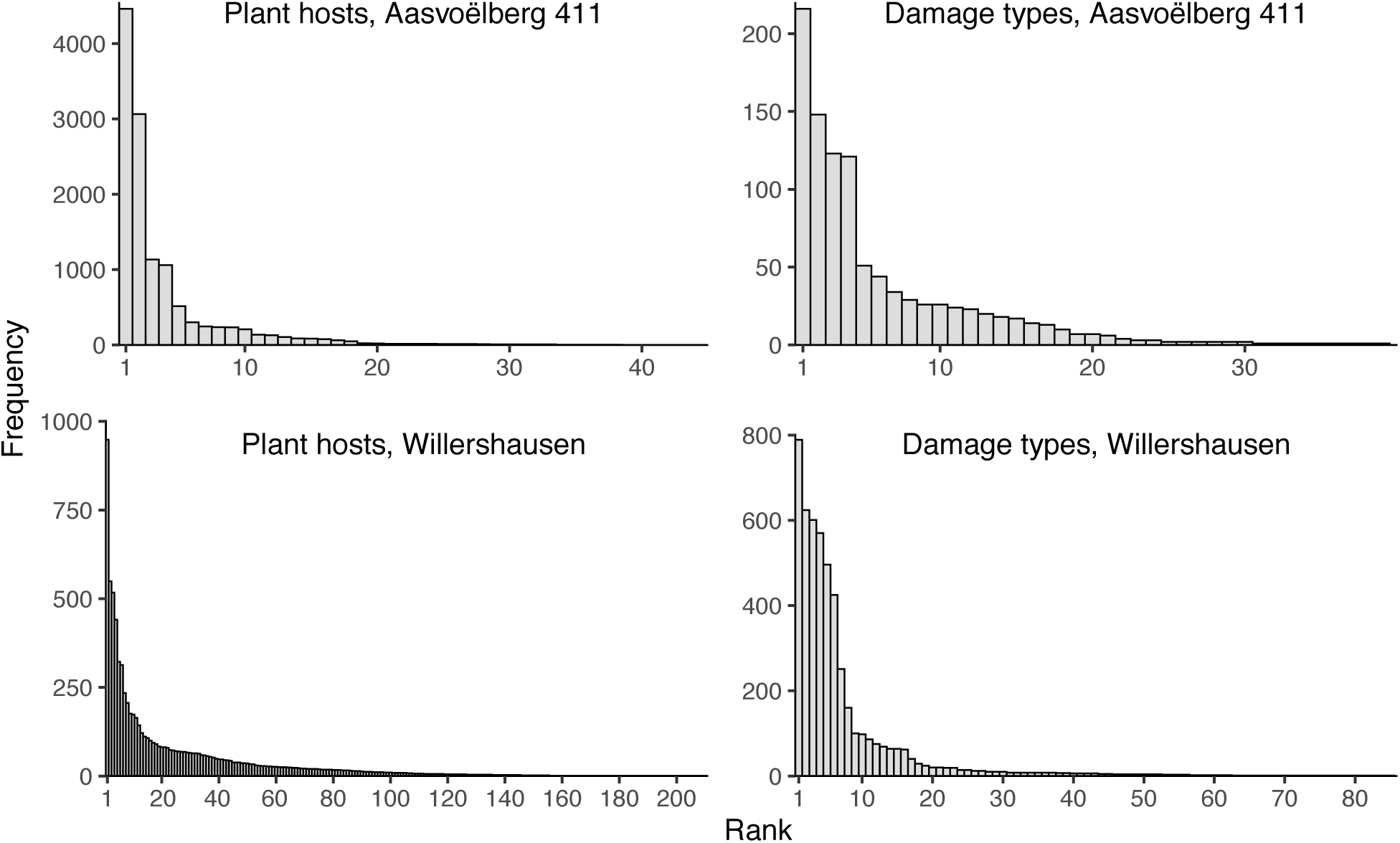
Dominance-diversity distributions for plant hosts and damage types at the pre-angiosperm assemblage with the largest number of leaves examined, Aasvoëlberg 411 (Labandeira et al., 2018), and the angiosperm assemblage with the largest number of leaves examined, Willershausen (Adroit et al., 2018).

**Figure S6:**
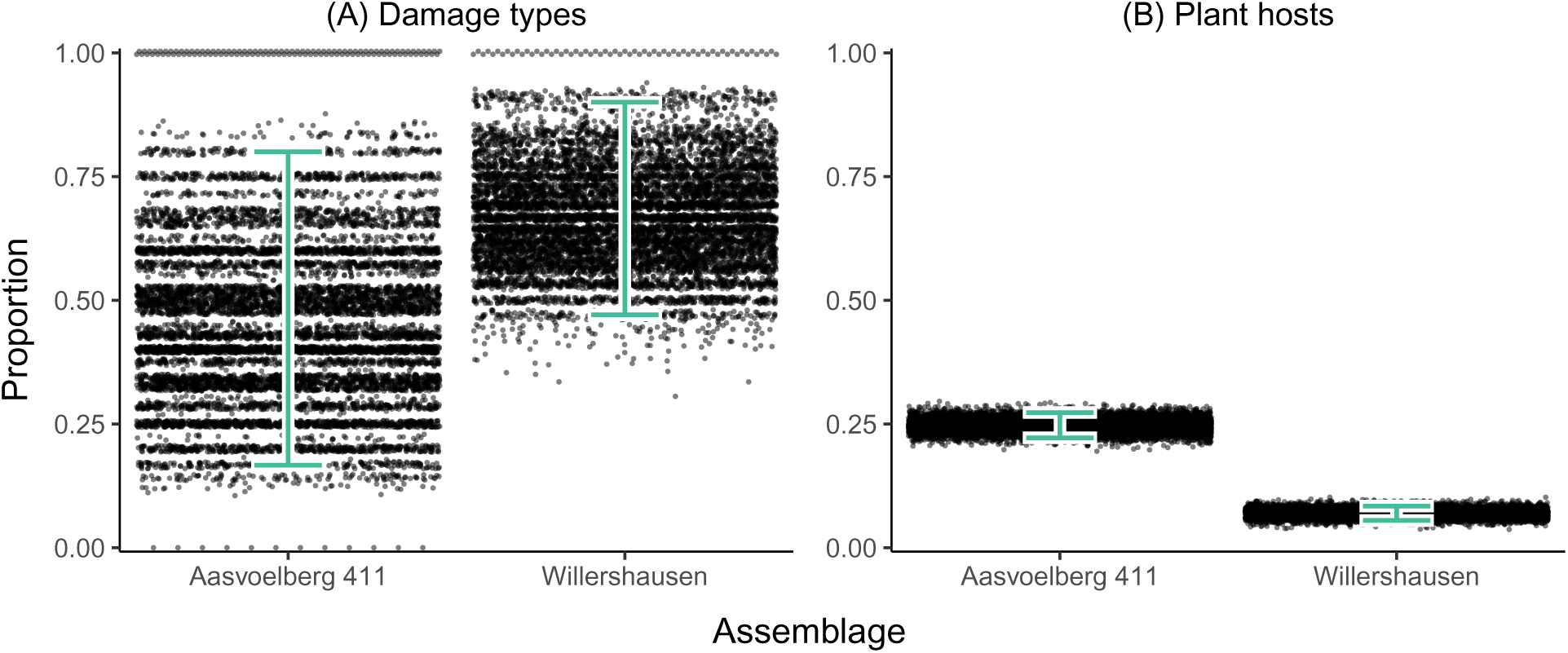
Measures of community structure at Aasvoëlberg 411 and Willershausen, the two assemblages featured in Figure S5, calculated by subsampling each assemblage to 1,000 leaves. Panel (a) shows the proportion of damage types on the second-most-abundant plant host that also occur on the most abundant plant host. Panel (b) shows the number of leaves belonging to the second-most-abundant plant host divided by the number of leaves belonging to the most abundant plant host. The lines denote 95% confidence intervals.

**Figure S7:**
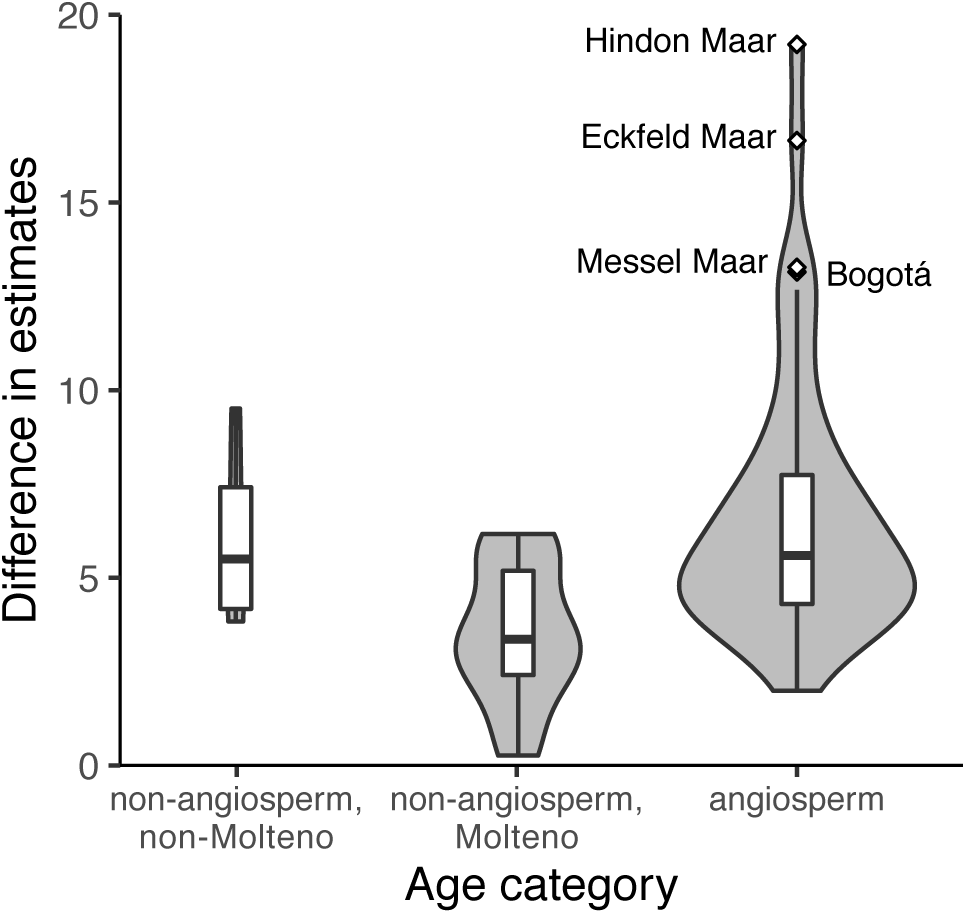
The difference in estimated damage type diversity when the sample coverage used in rarefaction increases from 0.8 to 0.9. The width of each violin represents the number of assemblages it contains. Boxplots are overlain atop each violin.

**Figure S8:**
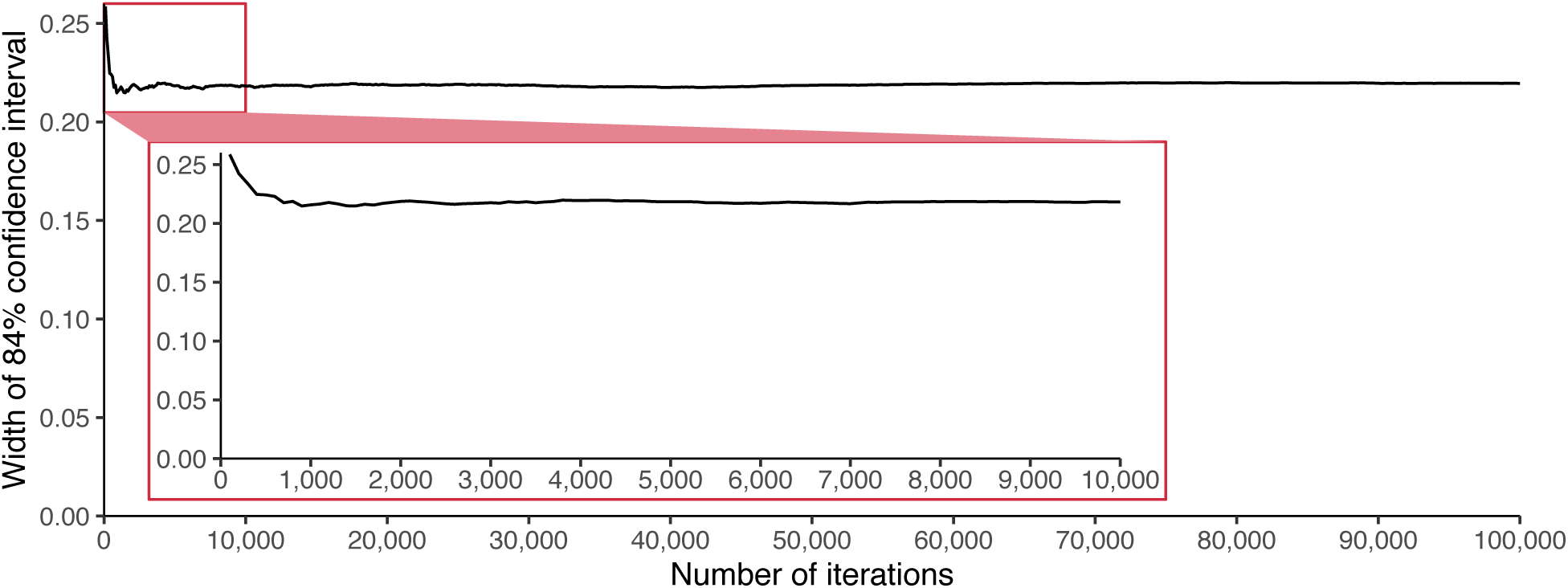
Changes in the width of the 84% confidence interval as the resampling procedure is further iterated.

